# Quantatitive Analysis of Conserved Sites on the SARS-CoV-2 Receptor-Binding Domain to Promote Development of Universal SARS-Like Coronavirus Vaccines

**DOI:** 10.1101/2021.04.10.439161

**Authors:** Siling Wang, Dinghui Wu, Hualong Xiong, Juan Wang, Zimin Tang, Zihao Chen, Yizhen Wang, Yali Zhang, Dong Ying, Xue Lin, Chang Liu, Shaoqi Guo, Weikun Tian, Yajie Lin, Xiaoping Zhang, Quan Yuan, Hai Yu, Tianying Zhang, Zizheng Zheng, Ningshao Xia

**Author notes:** These authors contributed equally (Siling Wang, Dinghui Wu, Hualong Xiong, Juan Wang). Correspondence and requests for materials should be addressed to: TY.Z, ZZ.Z, NS.X.

## Abstract

Although vaccines have been successfully developed and approved against SARS-CoV-2, it is still valuable to perform studies on conserved antigenic sites for preventing possible pandemic-risk of other SARS-like coronavirus in the future and prevalent SARS-CoV-2 variants. By antibodies obtained from convalescent COVID-19 individuals, receptor binding domain (RBD) were identified as immunodominant neutralizing domain that efficiently elicits neutralizing antibody response with on-going affinity mature. Moreover, we succeeded to define a quantitative antigenic map of neutralizing sites within SARS-CoV-2 RBD, and found that sites S2, S3 and S4 (new-found site) are conserved sites and determined as subimmunodominant sites, putatively due to their less accessibility than SARS-CoV-2 unique sites. P10-6G3, P07-4D10 and P05-6H7, respectively targeting S2, S3 and S4, are relatively rare antibodies that also potently neutralizes SARS-CoV, and the last mAbs performing neutralization without blocking S protein binding to receptor. Further, we have tried to design some RBDs to improve the immunogenicity of conserved sites. Our studies, focusing on conserved antigenic sites of SARS-CoV-2 and SARS-CoV, provide insights for promoting development of universal SARS-like coronavirus vaccines therefore enhancing our pandemic preparedness.

## Introduction

Severe acute respiratory syndrome coronavirus 2 (SARS-CoV-2) causes the ongoing outbreak of coronavirus disease 2019 (COVID-19), resulting in a global pandemic (*1, 2*). As of April 9, 2021, SARS-CoV-2 has caused more than 133 million people infection around the world including about 3 million deaths, which lead to unprecedented enormous global health and economic damage. The ~30 kb RNA genome of SARS-CoV-2 encodes four structural proteins including the spike (S), membrane (M), envelope (E) and nucleocapsid (N) proteins, and non-structural proteins, as well as a number of accessory proteins (*3*). The transmembrane spike glycoprotein is divided into S1 comprising receptor binding domain (RBD) and N-terminal domain (NTD), and S2 which promotes membrane fusion by fusion peptide. As most neutralizing mAbs isolated from convalescent COVID-19 individuals target RBD (*4–8*), by which S protein binds to receptor angiotensin-converting enzyme 2 (ACE2) and promotes exposure of fusion peptides within S2 component contributing to viral membrane fusion with host cells, it is main target for design of therapeutics and vaccine (*9, 10*).

Human coronaviruses (HCoVs) infecting human are composed of HCoV-OC43, HCoV-HKU1, HCoV-229E and HCoV-NL63, as well as the highly pathogenic MERS-CoV, SARS-CoV and SARS-CoV-2. SARS-CoV-2 phylogenetically close to SARS-CoV is classified into the *Betacoronavirus* genus including another highly pathogenic MERS-CoV, as well as HCoV-OC43 and HCoV-HKU1 variants leading to endemic (*11, 12*). Historically, there have been three HCoVs infections causing severe syndrome, including SARS outbreak that initially identified as an exotic infection in coronaviruses evolution, MERS-CoV outbreak as a second hard hit, and current COVID-19 pandemic (*13–16*). These studies reveal that it is reasonable to speculate on the possibility of emergence of other SARS-like coronavirus in the future. Hence, it is certainly valuable to promote development of more universal coronavirus vaccines and broader therapeutic agents by characterization of conserved antigenic sites in SARS-CoV-2 RBD to enhance our preparedness against the possible pandemic-risk of SARS-like coronavirus in the future. Moreover, as the duration of the SARS-CoV-2 pandemic extends, currently, there has been multiple variants emerging around the world. Although D614G mutation in the S protein significantly promotes corresponding variants infectivity in susceptible cells, this residue substitutions fails to cause immune escape (*17–21*). Conversely, many results support that the predominant SARS-CoV-2 variant B.1.351 in South Africa, with many crucial mutations in RBD including K417N, E484K and N501Y, could decrease therapeutic efficacy of neutralizing antibodies and even compromise protective efficacy of approved SARS-CoV-2 vaccines targeting the initial SARS-CoV-2 that emerged in 2019 (*20, 22–25*). Hence, universal SARS-like coronavirus vaccines based on the conserved antigenic sites in RBD also is potential in prevention against highly pathogenic SARS-CoV-2 variants that could escape from the established specific immune memory.

Although SARS-CoV-2 and SARS-CoV share 90% amino acid identity in S2 domain, SARS-CoV-2 RBD just shows lower 73% amino acid identity with SARS-CoV RBD (*12*), implying that there may be less conserved antigenic sites within RBD. Nevertheless, some highly conserved epitopes on the SARS-CoV RBD have been identified by mAbs CR3022 (*26*), S309 (*27*) and ADI-56046 (*26*), the cross-binding mAbs that were originally isolated from SARS patient, of which S309 and ADI-56046 could efficiently neutralize infection of SARS-CoV-2 and SARS-CoV. In addition, many human mAbs targeting SARS-CoV-2 S protein have been reported, which were isolated from convalescent COVID-19 individuals, however, cross-binding mAbs are relatively rarely shown, especially cross-neutralizing mAbs (*29–31*), indicating that SARS-CoV-2 conserved antigenic sites within RBD is subimmunodominant compared to its unique sites. While some conserved antigenic sites have been identified by cross-binding mAbs including CR3022 (*26*), S309 (*27*) and ADI-56046 (*28*) from SARS-CoV survivors as well as COVA1-16 (*5*), EY6A (*8*) and 2-36 (*32*) from COVID-19 individual, no studies are performed to quantitatively define antigenic map of conserved sites in SARS-CoV-2 RBD.

In this study, 77 SARS-CoV-2 RBD-specific antibodies were isolated from a cohort of 10 convalescent COVID-19 individuals for further biophysical characterization, by which we succeeded to define a quantitative antigenic map of neutralizing sites within SARS-CoV-2 RBD. We found that sites S2, S3 and S4 were conserved sites and were subimmunodominant compared with SARS-CoV-2 unique site S1. Unlike SAS-CoV-2 unique sites, conserved sites S2, S3 and S4 are proved as less accessible sites, weakly inducing antibody affinity mature. Moreover, to improve antibody response to conserved sites, we tried to design some RBDs, which contributed to development of universal SARS-like coronavirus vaccines.

## Results

### Seroconversion of antibodies against SARS-CoV-2 in convalescent COVID-19 individuals and isolation of SARS-CoV-2 RBD-specific antibodies

We collected blood samples from 10 convalescent COVID-19 individuals between February and March 2020, who were previously infected with SARS-CoV-2, confirmed by PCR (polymerase chain reaction), and accompanied with fever, cough and other symptoms (Table. S1). The humoral immune response was efficiently elicited for these convalescent individuals with high plasma titer of RBD-specific IgG with difference less than an order of magnitude (Table. S2). Since plasma neutralizing capacity strongly correlates with RBD-specific total antibody and IgG titer, indicating that RBD of S protein is dominant target of neutralizing antibodies elicited by SARS-CoV-2 infection, plasma titer of RBD-specific total antibody and IgG titer might be identified as effective evaluation indicator in the selection of COVID-19 convalescent volunteers for collection of potent neutralizing plasma in emergency circumstances (Fig. S1). The proportion of RBD-specific B cells in memory B cells ranged from 0.03% to 0.18%, and the RBD-specific memory B cells contained higher percentage of IgG subtype than IgM subtype, revealing that SARS-CoV-2 infection efficiently promoted B cell receptor (BCR) class switch and affinity maturate in convalescent individuals, thus eliciting strong humoral immune response against SARS-CoV-2 (Fig. S2A, B and C). Nevertheless, how SARS-CoV-2 RBD-specific antibodies inhibit infection is still not clear, to this end, 77 RBD-specific monoclonal antibodies (mAbs) were obtained from 10 convalescent individuals for comprehensive feature description (Fig. S2D).

### Characterization of SARS-CoV-2 RBD-specific mAbs obtained from convalescent COVID-19 individuals

To identify the SARS-CoV-2 RBD-specific mAbs repertoire usage, we compared sequence with well-defined naïve repertoire of IMGT-database to obtain assigned V region germline. Based on exclusion of clonal expansion just in P01, P03 and P04, 67 unique clonotypes were identified. Notably, it was found that 7 out of 8 antibody sequences obtained in P03 individual were highly conserved, except for P03-3B1, illustrating that this clone type BCR was immune dominant clone in P03, induced by SARS-CoV-2 infection (Fig. 1A and Fig. S3). Furthermore, enrichment of multiple VH and VK/VL, including VH1-69, VH3-13, VH3-30, VH3-53, VK1-39 and VL1-40, were observed, in addition, VK1-39 derived light chains were most often combined with heavy chain of various types VH to form antibodies, accounting for 22.4% (15/67) (Fig. S4 and Fig. 1A). The mean somatic hypermutation (SHM) rate of heavy chain V region was similar among individuals, and the mean levels (2%) are comparable with those detected in infection of other respiratory virus (Fig. 1B) (*33–35*). Additionally, even though the average length of CDRH3 of RBD-specific mAbs was consistent with that of naïve repertoire (~15 amino acids), we observed significant enrichment of shorter length CDRH3 (11 amino acids) in VH3-53/66 derived mAbs, differing from influenza virus and HIV-1 binding mAbs (Fig. 1C and Fig. S8A) (*36–38*).

**Fig. 1.**
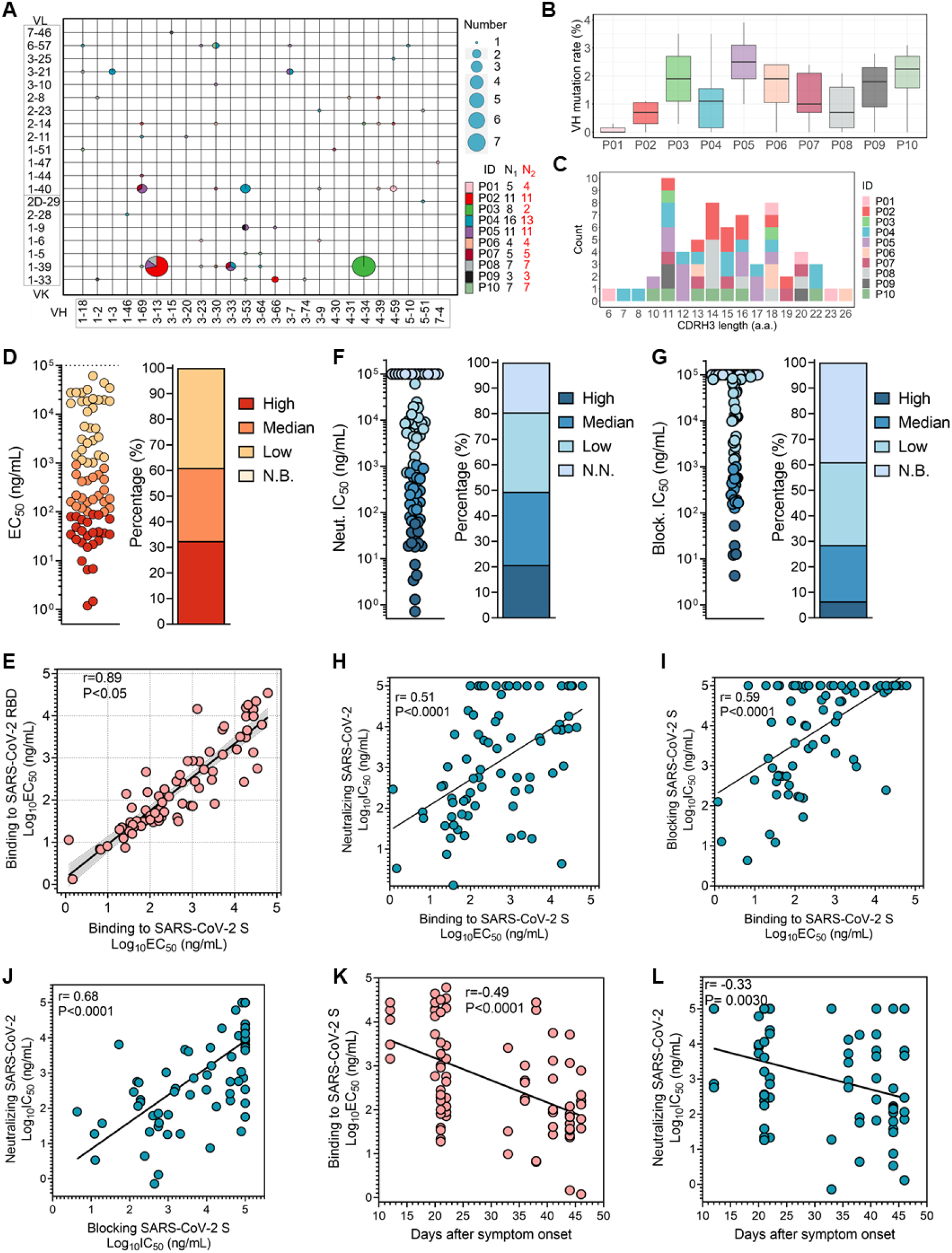
Characterization of SARS-CoV-2 RBD-specific mAbs obtained from convalescent COVID-19 individuals. **(A)** V gene frequencies for heavy and light chains of SARS-CoV-2 RBD-specific antibodies. The size corresponding to the number of heavy and light chain pairs in the repertoires is also denoted. Color indicates different convalescent individuals. N1 indicates number of SARS-CoV-2 RBD-specific antibodies for different individuals, and N2 indicates number of SARS-CoV-2 RBD-specific antibodies with unique clonotype for different individuals. **(B)** The V region SHM of heavy chain of specific antibodies from different individuals (N=67). **(C)** Distribution of CDR3 length of heavy chain. Antibodies are colored by each individual (N=67). V region germline and SHM and CDR3 length are determined using the Immunogenetics (IMGT). **(D)** The binding activity of specific antibodies to SARS-CoV-2 S protein are determined by ELISA. The color corresponds to binding activity, High (EC_50_ <100ng/mL), Median (EC_50_ between 100ng/mL and 1μg/mL), Low (EC_50_ between 1μg/mL and 100 μg/mL) and N.B. (EC_50_ >100 μg/mL). **(E)** Correlation between binding capacity to SARS-CoV-2 RBD and S protein (N=77). The 95% confidence interval of the regression line is shown in light grey, and r and P value of the correlation are also indicated. **(F and G)** Neutralizing capacity are determined by SARS-CoV-2 pseudovirus, and blocking capacity determined by S protein binding model. IC50s are showed in left panel, and percentage of mAbs with the indicated IC50 range is showed in right panel. The color represents different neutralizing capacity and blocking capacity, High (IC_50_ <100ng/mL), Median (IC_50_ between 100 ng/mL and 1 μg/mL), Low (IC_50_ between 1 μg/mL and 100 μg/mL), and N.N. or N.B. (IC_50_ >100 μg/mL). **(H and I)** The correlation between binding activity and neutralization or blocking capacity. r and P value of the correlation are also indicated. **(J)** The correlation between neutralization and blocking capacity, and r and P value of the correlation are indicated. **(K)** Correlation between binding activity of specific antibodies and duration of immune response for convalescent individuals (N=77). **(L)** The change of neutralization potency of specific mAbs during days after symptom onset. r and P value of the correlation are indicated. All correlation analysis is performed using Spearman test.

Subsequently, to determine profile of binding activity to SARS-CoV-2, we assessed binding activity of these mAbs to the recombinant S protein and RBD protein of SARS-CoV-2 using enzyme linked immunosorbent assay (ELISA). mAbs obtained from convalescent individuals presented binding activity to SARS-CoV-2 S protein in diversity, 61.0% of which showed strong binding activity (EC_50_ <1 μg/mL) to SARS-CoV-2 S protein, suggesting that infection of SARS-CoV-2 can effectively stimulate the humoral immune response producing a large number of high affinity specific mAbs (Fig. S2A and Fig. 1D). Interestingly, while there was correlation between binding activity to SARS-CoV-2 RBD and to SARS-CoV-2 S protein, some mAbs bound more strongly to RBD protein compared to S protein, implying that their targeting antigenic epitopes were poorly presented to be recognized in S protein, due to the coverage by other RBD monomer or NTD domain, as previously reported (Fig. 1E) (*39*). Next, we assessed neutralizing activity of these mAbs using a VSV pseudovirus model carrying SARS-CoV-2 S protein, and the neutralization IC_50_ potencies were shown in Fig. S5C. 80.5% (62/77) of these mAbs displayed neutralization against SARS-CoV-2, characterized as neutralizing antibodies (NAbs), among them 16 were identified as potent neutralizer with IC_50_ < 0.1 μg/mL, 22 as moderate neutralizer with IC_50_ of 0.1-1 μg/mL, and 24 as weak neutralizer with IC_50_ of 1-100 μg/mL (Fig. 1F). Surprisingly, none of the antibodies isolated from P03 convalescent individual had the potent neutralizing capacity, although amount of antibodies derived from the same clone were elicited (Fig. S6).

To investigate whether blocking S protein binding to ACE2 is the key mechanism for NAbs to inhibit infection of SARS-CoV-2 into susceptible cells, such as lung cells and intestinal cells, mAbs were mixed with SARS-CoV-2 S protein at different concentrations and then added into 293T cells expressing human ACE2 to evaluate the capability of blocking S protein entrance into cells, as reported (*40*). The results showed that 61.0% of the specific mAbs had the ability to block entrance of S protein into cells, IC_50_ ranging from 4 ng/mL to 100 μg/mL (Fig. S5D and Fig. 1G). These blocking and neutralizing capacity well correlated with the binding capacity (Fig. 1H and I). In terms of the neutralization and blocking data, we found their well correlation, indicating that blocking attachment of SARS-CoV-2 S protein to receptor ACE2 was critical to inhibit SARS-CoV-2 infection by NAbs targeting SARS-CoV-2 RBD (Fig. 1J and Fig. S7). But not all of them, some NAbs neutralized SARS-CoV-2 with weakly blocking SARS-CoV-2 S protein binding to ACE2, which is similar with previously observation (*27*). Although the corresponding neutralizing mechanism has not been explained clearly, it is putative that binding of these NAbs may impede sequential conformational changes of S protein for SARS-CoV-2 genome intracellular release.

Abnormally enrichment of mAbs with CDRH3 of 11 amino acids displayed strong binding activity to SARS-CoV-2 S protein, majority of them were encoded by VH3-53/66. While recent reports demonstrated that these mAbs have innate binding capability to SARS-CoV-2 RBD, with shorter CDRH3 for reduction of collision with RBD, length of CDRH3 did not correlate with binding activity (Fig. S8). To determine whether affinity of SARS-CoV-2 RBD-specific mAbs efficiently evolves, we analyzed the association between binding activity of mAbs and duration of immune response. The binding activity and neutralizing capacity of specific mAbs correlated with days after symptom onset, illustrating that affinity of these specific mAbs has the ability to continuously evolve (Fig. 1K and L and Fig. S9). However, plasma anti-RBD IgG titer did not correlate with days after symptom onset for individuals, putatively due to decay of corresponding specific mAbs production in blood (Fig. S1A) (*41*).

### Profile of cross-binding RBD-specific antibodies

Due to 76% sequence identity shared between SARS-CoV S protein and SARS-CoV-2 S protein, humoral immune could utilize some conserved epitopes to produce cross-reactive mAbs (*42*). As expected, some of SARS-CoV-2 specific mAbs also bound to SARS-CoV S protein, accounting for 50.6% of SARS-CoV-2 specific antibodies, of them 11 (14.3%) recognizing SARS-CoV S protein with strong binding activity (EC_50_ <1 μg/mL) (Fig. 2A and Fig. S5B). Expectedly, these cross-binding mAbs also showed continuously affinity maturation (Fig. 2B). It was observed that mAbs encoded by some VH germline had a tendency to broadly reactive with SARS-CoV and SARS-CoV-2, such as VH1-69, VH3-13, VH3-30 and VH4-46 (Fig. 2C). Additionally, cross-binding mAbs showed tendency of lower SHM in VH and JH region, which might limit their affinity maturation, putatively due to less exposure of corresponding conserved antigenic sites (Fig. 2D and E). Overall, these results confirmed efficient cross-binding antibody response during SARS-CoV-2 infection and the presence of conserved antigenic sites inducing the maturation of cross-reactive mAbs.

**Fig. 2.**
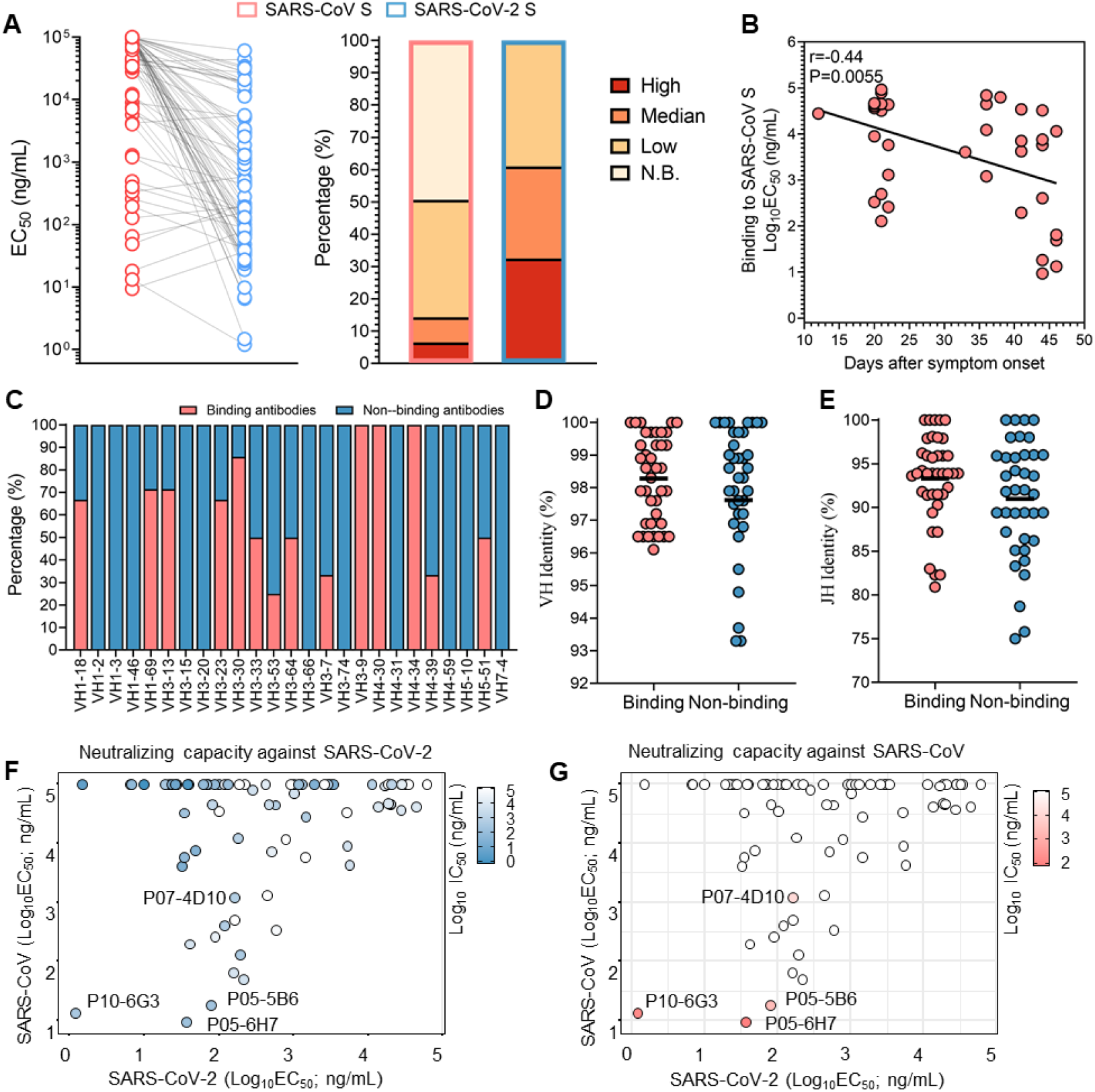
Profile of cross-binding SARS-CoV-2 RBD-specific mAbs. **(A)** The binding activity of specific antibodies to SARS-CoV S protein and SARS-CoV-2 S protein are determined by ELISA. The color corresponds to binding activity, High (EC_50_ <100 ng/mL), Median (EC_50_ between 100 ng/mL and 1 μg/mL), Low (EC_50_ between 1 μg/mL and 100 μg/mL) and N.B. (EC_50_ >100 μg/mL). **(B)** Correlation between binding activity to SARS-CoV S protein and duration of immune response. Correlation analysis is performed using Spearman test, and r and P value of the correlation are also indicated. **(C)** Distribution of SARS-CoV binding mAbs in VH region germline. **(D and E)** Comparison of VH and JH mutation between SARS-CoV S protein binding mAbs and non-SARS-CoV S protein binding mAbs. Black line denotes mean value. **(F and G)** Neutralizing capacity against SARS-CoV pseudovirus and SARS-CoV-2 pseudovirus are shown in the context of binding activity. The color varies with the neutralizing capability, deep blue and deep red for strong neutralizing capability.

Notably, the majority of mAbs with potent neutralization against SARS-CoV-2 showed no reactivity with SARS-CoV, confirming that unique antigenic sites in SARS-CoV-2 RBD, as immunodominant epitopes, more efficiently elicit high affinity mAbs with potent neutralizing capacity than conserved epitopes (Fig. 2F). Subsequently, cross-binding mAbs were tested for their ability to neutralize SARS-CoV. Our results indicate that only P10-6G3, P07-4D10, P05-6H7 and P05-5B6 were identified as cross-neutralizing mAbs against SARS-CoV and SARS-CoV-2, of which P10-6G3 displayed more highly binding activity to SARS-CoV-2 S protein than SARS-CoV S protein, revealing that the SARS-CoV-2 unique residues of corresponding conserved site might be more accessible (Fig. 2F and G). Combined with binding activity, we found that the low affinity with SARS-CoV S protein should be the reason why a large number of cross-binding antibodies could not neutralize SARS-CoV (Fig. 2G). As the antigenic sites recognized by these cross-neutralizing mAbs were common between SARS-CoV and SARS-CoV-2, they can become the key targets for design of universal vaccine against SARS-like coronavirus and for selection of broadly therapeutic antibodies.

### Functional characterization of NAbs recognizing multiple different neutralizing antigenic sites

To investigate key neutralizing antigenic sites eliciting potent immune response, first of all, competition-binding assay using ELISA were performed for neutralizing mAbs. Followed by cluster analysis, the neutralizing antibodies were classified into 6 clusters (C1-6), according to the competitive ability resulting from sterical incompatibility when simultaneously binding to the same SARS-CoV-2 RBD antigenic sites, and the antigenic sites for C1-6 NAbs were respectively termed as sites S1-6 (Fig. S10 and S11 and Fig. 3A). Sites S1, S3 and S4 should be immunodominant antigenic sites, because of C1, C3 and C4 NAbs accounting for higher proportion (Fig. 3A). Notably, these NAbs of different clusters were induced in many SARS-CoV-2 convalescent individuals, which embodied common immunogenic characteristics of antigenic sites in RBD protein and indicated that infection of SARS-CoV-2 can elicit similar specific antibody response (Fig. 3B).

**Fig. 3.**
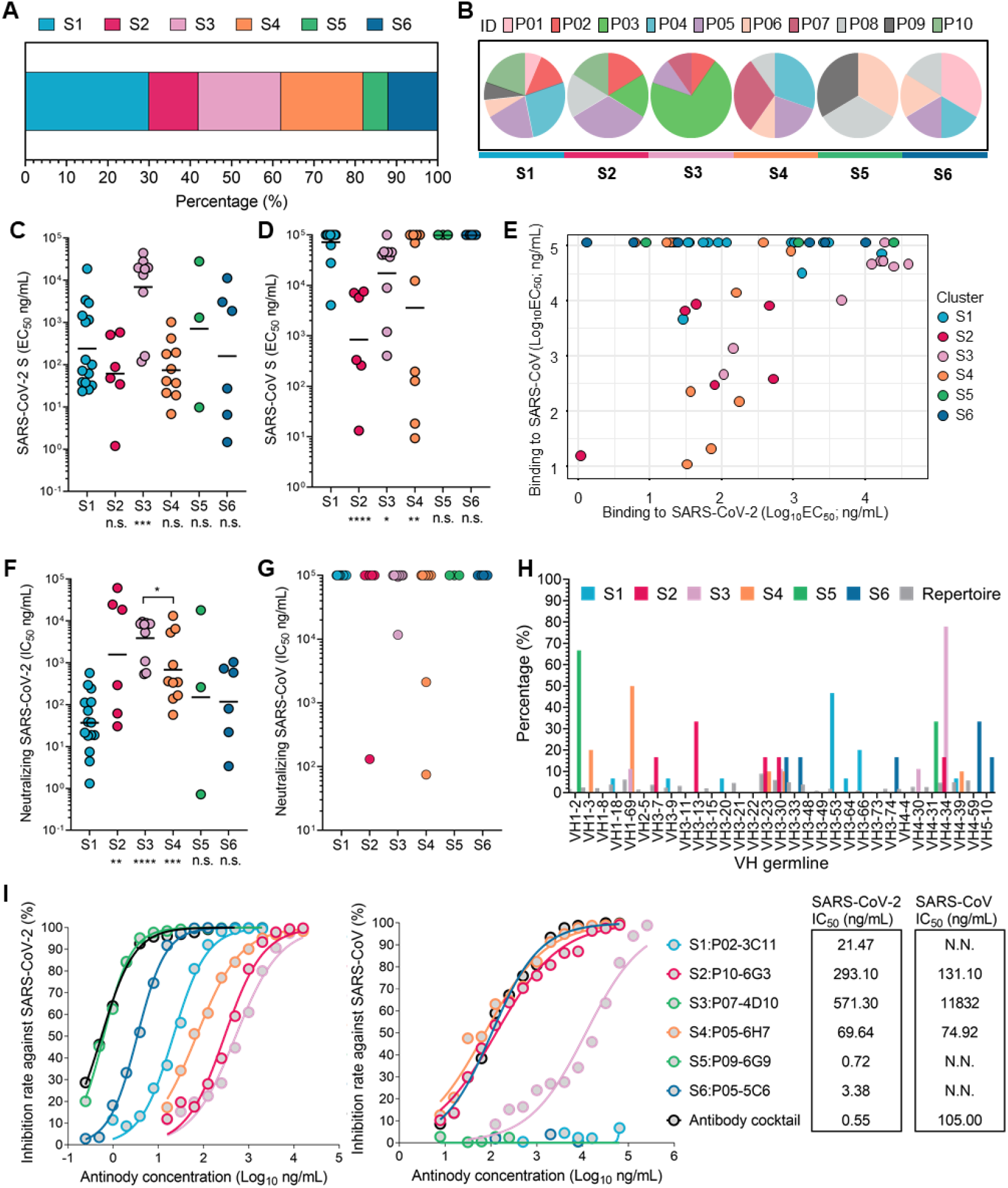
Multiple neutralizing epitope mapping of mAbs by clustering analysis and functional characterization. **(A)** Depending on clustering analysis of competition ELISA data for neutralizing antibodies, NAbs are classified into 6 clusters targeting 6 different RBD antigenic sites (S1-6). The percentage of NAbs recognizing different antigenic sites is calculated and displayed. Colors indicate different individuals. **(B)** Individual composition analysis of NAbs targeting different antigenic sites. ID denotes different convalescent individuals marked by colors. **(C and D)** The binding activity to SARS-CoV and SARS-CoV-2 S protein are determined by ELISA, and denoted by EC_50_. **(D)** Analysis of antigenic sites recognized by cross-binding NAbs. **(F and G)** Neutralizing capacity against SARS-CoV-2 pseudovirus (F) and against SARS-CoV pseudovirus (G) of NAbs targeting S2-6 are compared with S1 directed NAbs. **(H)** VH germline of each cluster neutralizing antibodies is analyzed. Different color indicates each neutralizing cluster. **(I)** Neutralization capacity of combination of representative NAbs targeting sites S1-6 against SARS-CoV-2 pseudovirus and SARS-CoV-2 pseudovirus. For panel C, D, F and G, data were plotted as the geometric mean. n.s.: no significant difference; *: P < 0.05; **: P< 0.01; ***: P < 0.001. The black line indicates mean value. Statistical significance in C, D, F and G using Mann-Whitney U test.

To determine the functional characteristics of NAbs targeting different sites, we analyzed the biochemical properties of them elicited by sites S1-6. NAbs of C1, C2, C4, C5 and C6 bound to SARS-CoV-2 S protein with comparable EC_50_, however, NAbs elicited by site S3 displayed lower binding activity to SARS-CoV-2 S protein (Fig. 3C). Furthermore, majority of C2, C3 and C4 NAbs showed cross-reactivity with SARS-CoV S protein and SARS-CoV-2 S protein, illustrating that these sites were conserved sites between SARS-CoV and SARS-CoV-2 (Fig. 3D and E). Nevertheless, these conserved sites just induced 4 potent cross-neutralizing mAbs (P10-6G3 targeting S2; P07-4D10 targeting S3; P05-5B6 and P05-6H7 targeting S4) against SARS-CoV and SARS-CoV-2 in natural infection of SARS-CoV-2, since most cross-binding NAbs showed obviously weaker affinity with SARS-CoV S protein compared to SARS-CoV-2 S protein, leading their incapacitation of cross-neutralization (Fig. 3C, E, F and G). Notably, except conserved site S3, the remaining conserved sites directed NAbs showed lower neutralization capacity against SARS-CoV-2 than NAbs recognizing SARS-CoV-2 unique antigenic sites (sites S1, S5 and S6), under the premise of possessing similar binding activity to SARS-CoV-2 S protein, implying that these NAbs targeting conserved sites have a disadvantage in blocking S protein binding to receptor, putatively due to their partial overlap with ACE2 site (Fig. 3C and F).

C1 NAbs isolated from overwhelming majority of convalescent individuals recognized site S1 and potently specifically inhibited SARS-CoV-2 infection by blocking S protein attachment to ACE2, revealing that site S1 is immunodominant antigenic site in SARS-CoV-2 RBD to efficiently elicit strong NAbs response during SARS-CoV-2 natural infection (Fig. 3F and Fig. S12A). The immunodominance of site S1 may result from either its accessibility in different conformation of SARS-CoV-2 S protein or the innate affinity of corresponding C1 NAbs derived from naïve B repertoire (such as VH 3-53/66 germline), the latter factor will lead to the rapid affinity mature of C1 mAbs without the need for high level of somatic hypermutation (*31, 43, 44*) (Fig. 3H). On the contrast, while large amplification of the same antibody clone was exhibited against site S3 in convalescent P03, the NAbs of such antibody clone poorly inhibited infection of SARS-CoV-2, revealing that the site S3 is possibly immunodominant weakly-neutralizing site (Fig. 3F, Fig. S6, Fig. 1A and Fig. S4). In addition, site S4 elicited many neutralizers with weak blocking capacity, such as P05-6H7 and P06-6E10, these NAbs might inhibit SARS-CoV-2 infection either by inhibiting conformational change of S protein necessary for membrane fusion or by enhancing the S1 domain shedding to block SARS-CoV-2 attachment to ACE2 (Fig. 3F, G and Fig. S12). We further selected representative antibodies targeting sites S1-6 (C1, C2, C3, C4, C5 and C6 cluster represented by P02-3C11, P10-6G3, P07-4D10, P05-6H7, P09-6G9 and P05-5C6, respectively), showing potently neutralizing capacity, as potential compatible therapeutics against COVID-19. The antibody cocktail demonstrated potently and broadly neutralizing capacity across SARS-CoV and SARS-CoV-2, determined by previously reported pseudovirus neutralization assay (Fig. 3I) (*45*). These mAbs recognized different SARS-CoV-2 RBD sites so as to avoid immune escape caused by virus mutation. Taken together, sites S2, S3 and S4 were identified as conserved antigenic sites in SARS-CoV-2 RBD that could induce cross-neutralizing mAbs response and are considerable in rational design of universal SARS-like coronavirus vaccines, and sites S1, S5 and S6 as SARS-CoV-2 unique antigenic sites. Difference in binding location possibly contributes to that NAbs elicited by conserved antigenic sites showed weaker neutralization against SARS-CoV-2 than those by unique antigenic sites.

### Identification of antigenic sites S1-6 on SARS-CoV-2 S protein

Characterization of key antigenic sites conformation is essential to present a great promise for designing novel vaccine against SARS-CoV-2, and the conserved antigenic sites remain to be further studied for rational design of universal vaccine against SARS-like coronavirus. To determine the spatial position of site S1-6, we performed mutagenesis study of SARS-CoV-2 RBD by substituting ACE2-interactive and non-interactive residues with alanine or arginine, and then assessed the decreased binding activity of representative NAbs to mutant RBDs compared to the reference RBD, as previously reported (*46*). The reduction of binding activity to each mutant RBDs relative to reference RBD were shown in Fig. S13. The spatial position of all antigenic sites was simultaneously displayed on RBD to demonstrate their relative location, which is critical to expound functional characteristics of all neutralizing antigenic sites in RBD (Fig. S14 and Fig. 4A). The sites S1, S5 and S6 largely overlapped with the binding site of ACE2, supporting potent neutralizing activities of corresponding NAbs by efficiently blocking S protein binding to receptor, whereas site S2, S3 and S4 were positioned away from ACE2 footprint (Fig. S14A). Hence, NAbs targeting site S3 inhibited infection with weak neutralizing capacity, likely due to poor steric hindrance with receptor ACE2. However, site S4-specific NAbs might neutralize infection of SARS-CoV-2 and SARS-CoV by novel mechanism without blocking S protein attachment to ACE2.

**Fig. 4.**
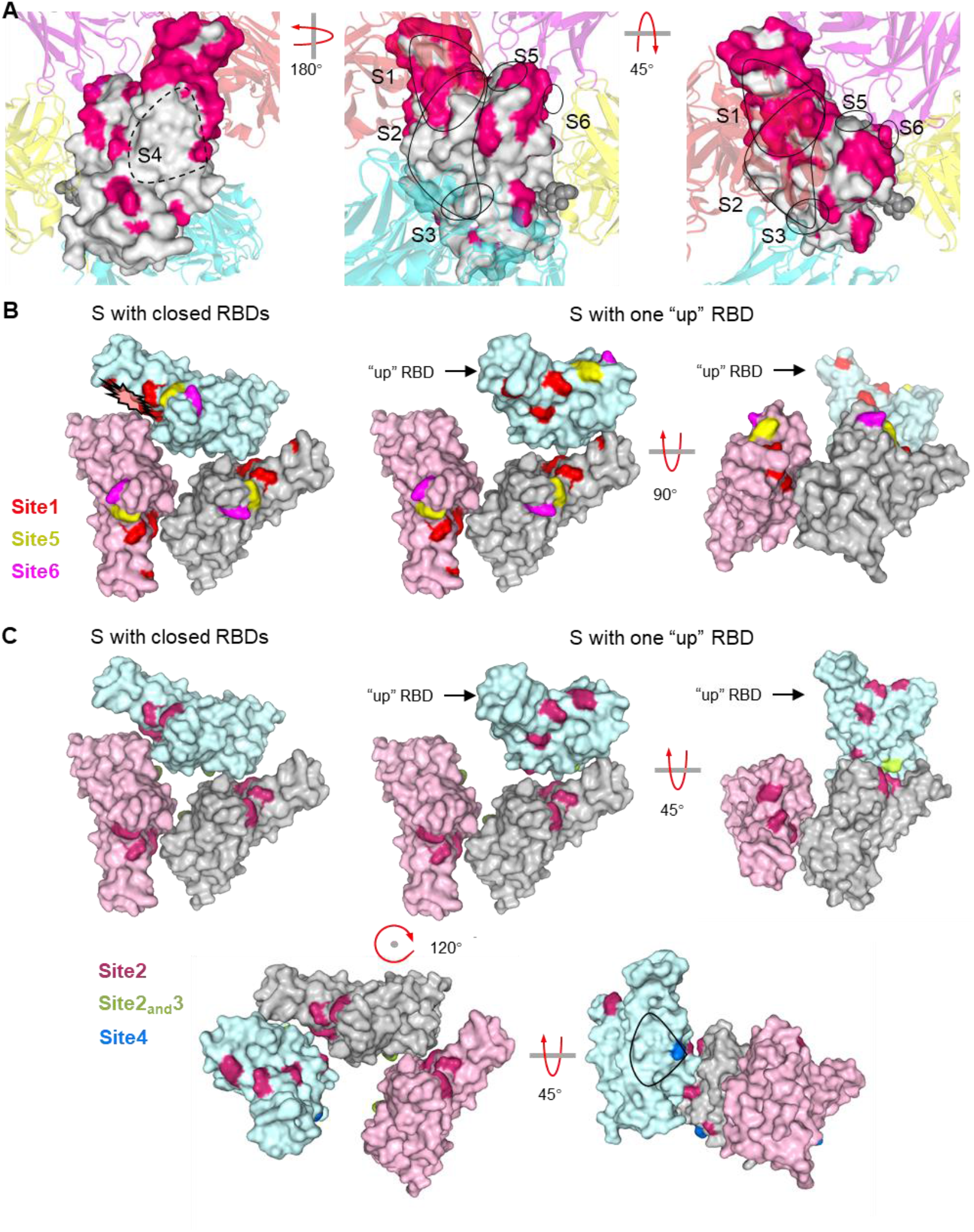
Dynamics analysis of sites S1-6 accessibility. **(A)** Relative position of sites S1-6 are shown on RBD. Some SARS-CoV-2 RBD-specific mAbs reported in other studies are displayed by different colors, and purple, red, cyan and yellow respectively indicate mAbs C102, P2B-2F6, S309 and S2A4, these mAbs are identified as representative mAbs for Class 1-4. Deep wine indicates different residues within RBD between SARS-CoV-2 and SARS-CoV. The glycan at position N343 is rendered as black spheres. **(B)** Accessible analysis of site 1, site5 and site 6 in either standing-up RBD or lying-down RBD of S protein. Red, yellow and purple denote critical residue of site S1, site S5 and site S6 on RBD of S protein, respectively (PDB code: 6VYB and 6VXX). **(C)** Accessible analysis of conserved sites S2, S3 and S4 in either lying-down RBD of S protein (PDB code: 6VXX) or standing-up RBD of S protein (PDB code: 6VYB). Red, yellow and purple denote critical residue of sites S2, S3 and S4.

S2, S3 and S4 were identified as conserved antigenic sites on RBD, due to the cross-reactivity of their corresponding mAbs, which was further confirmed by their critical residues, common between SARS-CoV and SARS-CoV-2 (Fig. 4A and Fig. S14B). Additionally, by summarizing results of SARS-CoV-2 research, it was found that the RBD neutralizing antigenic sites identified by us are similar to those by other studies (Fig. 4A) (*4, 47, 48*). Obviously, site S1 overlapped with Class 1 epitope, and site S5 and site S6 were adjacent to Class 2 epitopes recognized by representative C002 mAb. Site S2 was located between Class 1 epitope and Class 4 epitope, which might be similar to site IIa as previously reported (*4*). Up to now, epitopes like site S4 have not been reported, although S309 recognizing Class 3 epitope also showed similar functional property, neutralizing capacity without blocking S protein binding to ACE2 (*47*). As some neutralizing antigenic sites are hidden by RBD lying down state of S protein, it becomes important immune escape mechanism for SARS-CoV-2 (*39*). It was obvious that site S5 and S6 were accessible in either RBD lying-down state or RBD standing-up state, however, just RBD standing-up state of S protein made site S1 more accessible for corresponding NAbs (Fig. 4B). The conserved sites are highly concealed on lying-down RBD by adjacent RBD monomer, especially site S3. Even though RBD was in standing-up state, on which site S3 will not be sufficiently exposed, therefor, this inadequate exposure failed to improve affinity maturation of S3 directed mAbs (Fig. 4C, Fig. 3D and 3E). Similarly, no enough space for NAbs binding to conserved site S4 was found on closed S protein, and only RBD in standing-up state could improve the accessibility of site S4 (Fig. 4C). In summary, these results elucidate that the conserved antigenic sites displayed less accessibility than SARS-CoV-2 unique epitopes, and the poor accessibility have hindered affinity maturation of site S3 directing mAbs in natural infection and even will decrease the cross-neutralizing antibody response after SARS-CoV-2 vaccination.

### Rational design of SARS-CoV-2 RBD to enhance immunodominance of conserved antigenic sites

Majority of mAbs recognizing antigenic site S1 were VH3-53/66 mAbs with short (mostly 11 residues) CDRH3s. Moreover, many studies also have reported the same enrichment of VH3-53/66 mAbs targeting the antigenic sites similar to site S1, and find that they share structural similarities with each other by solving structures of immune complex with SARS-CoV-2 RBD (*8, 29, 48–50*). These VH3-53/66 mAbs showed native binding to SARS-CoV-2 RBD residues using CDRH1 and CDRH2 by forming many hydrogen bonds (L455, Y473, A475 and N487 bound by CDRH1, D420, Y421 and R457 bound by CDRH2) (Fig. 5A and Fig. S15). Therefore, the native binding advantage of antigenic site S1 to VH3-53/66 was the cardinal cause of its immunodominance, which can result in the massive amplification of antigenic site S1-specific B cells, while competing to inhibit the proliferation of B cells directing conserved antigenic sites. To indirectly enhance competitiveness of conserved antigenic sites for immune response, it is essential to decrease the immunodominance of site S1. To this end, we designed a variety of SARS-CoV-2 RBD proteins with antigenic site S1 silence using either protein truncation or glycan modification (Fig. 5B–5E). The glycan modification at position K458 and A475 within site S1, termed as RBDGlycan458,475, was successful to destroy the binding of S1 directed mAbs, including p02-3C11 derived from VH3-66 and P05-5C4 derived from VH3-53, and to maintain the remaining antigenic sites including conserved antigenic sites S2, S3 and S4, which might efficiently enhance the competitiveness of subimmunodominant conserved antigenic sites in immune response (Fig. 5F). To further investigate whether the RBDGlycan458,475 could improve cross-binding antibodies response, BALB/c mice were immunized with 20 μg/dose of RBD_Glycan458,475_. Two weeks after immunization, mice receiving RBD_Glycan458,475_ and reference RBD protein all presented detectable serum anti-SARS-CoV S IgG and anti-SARS-CoV-2 S IgG, however, RBD_Glycan458,475_ induced significantly higher cross-binding IgG titer compared to reference RBD (Fig. 5G). Hence, universal vaccines based on such glycan modification in SARS-CoV-2 RBD possibly induce stronger cross-binding antibody response that could efficiently protect from infection of SARS-like coronavirus and prevalent SARS-CoV-2 variants.

**Fig. 5.**
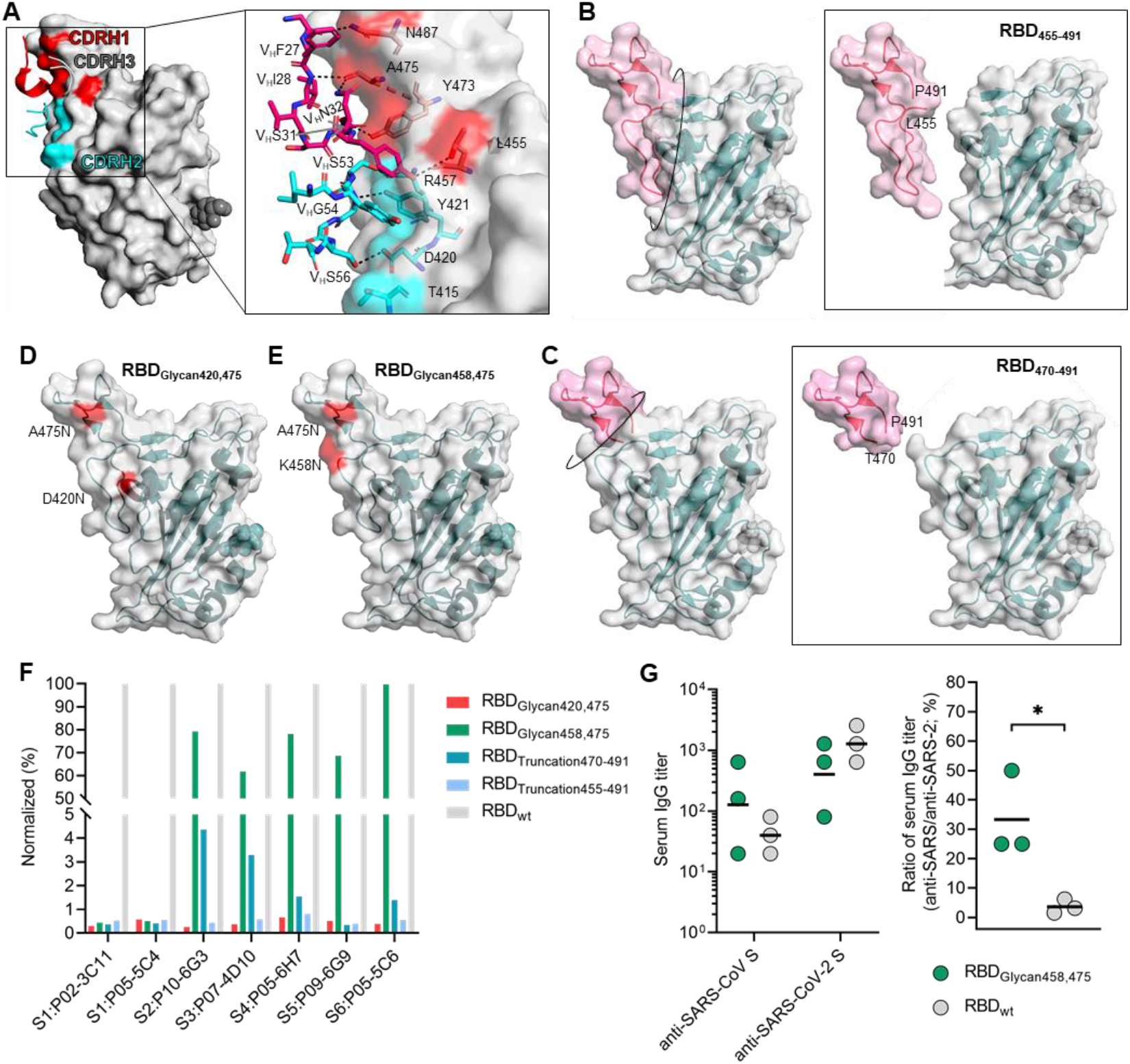
Structure of SARS-CoV-2 RBD site interacting with CHRH1 and CDRH2 of IGHV3-53/66 antibodies. **(A)** Interaction between CDRH1&2 residues and SARS-CoV-2 RBD. Residues of mAbs CDRH1&2 constructing hydrogen bond with RBD are rendered as red&cyan sticks. **(B, C, D and E)** Design of SARS-COV-2 RBD enhancing capability to elicit cross-reactive antibodies. Truncated SARS-CoV-2 RBD by cutting off T455-P491 are termed as RBD_455-491_ in **B**, and RBD by cutting off T470-P491 are termed as RBD_470-491_ in **C**. SARS-CoV-2 RBD glycosylated at position 420 and 475 are termed as RBD_Glycan420,475_ in **D,** and at position 458 and 475 are termed as RBD_Glycan458,475_ in **E**. **(F)** Influence of designed RBD on binding activity of representative NAbs targeting different antigenic sites. **(G)** Antibody response against SARS-CoV S protein and SARS-CoV-2 S protein induced by designed RBD_Glycan458,475_ in mice (N=3). BALB/c mice were immunized with 20 μg/dose at weeks 0, and serum specific IgG titers were tested at weeks 2. The ratio of serum IgG titer against SARS-CoV S protein and SARS-CoV-2 S protein was also calculated. For panel G, data were plotted as the geometric mean or mean. Statistical significance in G using non-paired t test, and * indicates P < 0.05.

## Discussion

This study provided a qualitative and quantitative analysis of SARS-CoV-2 RBD-specific antibody response through 10 convalescent COVID-19 individuals recruited between February and March 2020. SARS-CoV-2 RBD was defined as the immunodominant domain within SARS-CoV-2 S protein, as previously reported (*4*), putatively supported by its few glycosylation sites compared with the remaining S protein domains and higher accessibility on S protein with variable conformations, as well as by S1 domain shedding (*27, 28, 51*). In addition, 80.5% of RBD-specific mAbs displayed neutralizing capacity, of which 16 mAbs were identified as potent neutralizer with IC_50_ < 0.1 μg/mL further confirming RBD as immunodominant neutralizing domain, which results from not only its strong immunogenicity but interaction with receptor ACE2. Similar to others virus infection, an increase in mAbs binding activity to SARS-CoV-2 S protein accompanied by stronger neutralizing capacity over time was determined, related to ongoing affinity mature of antibody response, revealing that SARS-CoV-2 RBD-specific mAbs could maintain on-going affinity maturation. Interestingly, these SARS-CoV-2 RBD-specific mAbs obtained from convalescent individuals showed low level of somatic hypermutation, especially mAbs from P01, revealing that naïve BCR repertoire may contribute to rapid mAbs response to SARS-CoV-2 RBD and that some mAbs encoded by germline genes may have the advantages of innate ability to recognize SARS-CoV-2 RBD, such as mAbs derived from VH3-53/66 (*31, 43, 44*). These findings suggest that common neutralizing mAbs response might be efficiently induced by SARS-CoV-2 vaccine composed of RBD.

Structural studies have proved that the S protein possesses conformational dynamics, in which different pre-fusion conformations expose a variety of crucial epitopes, including conserved antigenic sites between SARS-CoV and SARS-CoV-2 that can be recognized by cross-binding mAbs (*31, 52*). Although some conserved antigenic sites, identified by cross-binding mAbs, have been reported, systematic analysis of conserved antigenic sites is still lacking, which is prejudicial to rational design of universal vaccine (*4–8*). We used information obtained from neutralizing mAbs to develop a quantitative antigenic map of SARS-CoV-2 RBD neutralizing sites that demonstrates immunodominance, neutralization properties and conserved properties. We found that site S2, S3 and S4 (these sites defined by mAbs P10-6G3, P07-4D10 and P05-6H7, respectively) are conserved antigenic sites that can elicit cross-binding mAbs response against SARS-CoV and SARS-CoV-2. However, these conserved antigenic sites are subimmunodominant, and S3 only induce lower-affinity mAbs response compared to site S1, an unique antigenic site that is immunodominant and coincides with the ACE2 footprint, putatively related to their lower accessibility in a variety of conformations. Even though adjacent RBD is in standing-up state, the site S3 and S4 will not be sufficiently exposed for specific-mAbs recognition. Additionally, the unique antigenic site S1 with the native binding advantage to VH3-53/66 might further suppress humoral immune response to the conserved antigenic sites through the depletion of a large number of B cells. mAbs P10-6G3, P07-4D10, P05-5B6 and P05-6H7 recognizing sites S2, S3 and S4 efficiently inhibit SARS-CoV and SARS-CoV-2 infection into vulnerable host cells, further confirming that these conserved sites could serve as crucial antigenic sites for universal vaccine. Although site S4 is far from ACE2 binding site, corresponding mAbs P05-5B6 and P05-6H7 can neutralize SARS-CoV and SARS-CoV-2 infection without blocking S protein binding to ACE2, as reported S309 (*47*). Majority of potently neutralizing antibodies inhibit viral infection by blocking spike proteins attachment with receptors (*53–55*), however, we speculate that the site S4 directed mAbs may inhibit SARS-CoV-2 and SARS-CoV infection by intracellular neutralization pathway, a mechanism inhibiting the release of virus within endosome into cytoplasm, as reported in recent studies (*40*).

Since SARS-CoV-2 unique antigenic sites (S1, S5 and S6) in RBD efficiently provoke specific antibody response and strongly inhibit the production of cross-binding antibodies, these antigenic sites, especially site S1, should be silenced for universal vaccine design. Moreover, some predominant SARS-CoV-2 variants B.1.351 and P.1 with K417N, E484K and N501Y mutations causing changes in antigenic sites, similar to unique antigenic sites S1, S5 and S6, promote evasion of antibody-mediated immunity obtained from natural infection or vaccination, however, no cross-binding mAbs displaying decreased binding activity to these variants have been reported (*20, 22–24*). These findings prove that it is difficult for universal vaccine based on unique antigenic sites in RBD to induce conserved antibodies response to prevent the possible pandemic-risk of SARS-like coronavirus in the future or persistent SARS-CoV-2 variants with antigenic drift, nevertheless, focusing on the conserved antigenic sites might have great potential for universal SARS-like coronavirus vaccines (*56*).

Similar strategies for universal vaccine design have been proposed for development of universal influenza virus vaccines that protect from infection with drifted seasonal and novel pandemic influenza virus strains (*57, 58*). Candidates of universal influenza vaccines are mainly based on the conserved antigenic sites in stalk domain of the hemagglutinin, for example, headless hemagglutinin structures and the display of conserved stalk-epitopes on nanoparticles, which currently shows promising results in animal models and has great reference significance (*59–62*). We defined site S1 as SARS-CoV-2 unique immunodominant site, the dominance is related to its greater accessibility compared to conserved sites, and the innate binding capability of corresponding mAbs derived from naïve B repertoire (VH 3-53/66 germline) using CDRH1 and CDRH2 (*63*). To indirectly enhance competitiveness of conserved sites for mAbs response by decreasing immunodominance of site S1, we designed a variety of SARS-CoV-2 RBD protein with site S1 silence by either removal of peptide fragment or glycan modification. In our study, the designed RBDGlycan458,475 with glycan modification destroying site S1 is successful to damage the binding of corresponding mAbs and to maintain the remaining sites including conserved sites S2, S3 and S4. By immunized using mice, RBDGlycan458,475 efficiently enhanced the competitiveness of subimmunodominant conserved sites to reach stronger cross-binding antibody response than reference RBD, revealing that such direction of SARS-CoV-2 RBD design could promote development of universal vaccines against SARS-like coronavirus.

In summary, our studies defined quantitative antigenic map of neutralizing sites within SARS-CoV-2 RBD and completed the characterization of conserved antigenic sites, which is requisite for rational design of universal vaccine. Moreover, we have tried to design some RBD proteins, aiming to enhance the immune competitiveness of conserved antigenic sites. Although SARS-CoV-2 vaccine has been developed and approved, our long lasting efforts aimed to prepare universal vaccine for other human epidemic of SARS-like coronavirus possibly prevalent in the future is still necessary.

## Acknowledgements

This work was supported by the National Natural Science Foundation of China [grant number 81993149041, 81702006], Science and Technology Major Project of the Fujian Province [grant number 2020YZ014001], Xiamen Science and Technology Major Project [grant number 3502Z2020YJ02] and National Key Research and Development Program of China [grant number 2016YFD0500301].

## Ethics approval

This study was approved by the Medical ethics committee of School of Public Health of Xiamen University.

