## Supplementary material for "Quantatitive Analysis of Conserved Sites on the SARS-CoV-2 Receptor-Binding Domain to Promote Development of Universal SARS-Like Coronavirus Vaccines": Fig. S1

### Supplementary Data

#### Materials and methods

##### Collection of blood samples

In this study a total of 10 COVID-19 convalescent individuals, infected with SARS-CoV-2 in January to March 2020, were enrolled. Their peripheral blood was collected after they were discharged from the first hospital of Xiamen University, and the results of virus nucleic acid test in the serum were negative. The study was approved by the institutional review board of the School of Public Health in accordance with the Declaration of Helsinki, and written informed consent was obtained.

##### Detection of plasma antibody titer against SARS-CoV-2 RBD

To detect the plasma titers of total antibodies (Ab), IgG and IgM against SARS-CoV-2 RBD, we performed commercial enzyme-linked immunosorbent assay (ELISA) kits (Beijing Wantai Biological Pharmacy Enterprise), according to the manufacturer's instructions. The Ab-ELISA kit is based recombinant viral antigen using a double-sandwich reaction form. The IgG-ELISA kit is an indirect ELISA assay, and the IgM-ELISA kit is based on μ-chain capture method. The samples were initially tested undiluted, and the positive samples with the signal to a cutoff ratio (S/CO) >=10 were further diluted (1:10, 1:100, 1:1,000 and 1:10,000) by PBS buffer containing 20% newborn bovine serum (NBS) and tested again. The titers for Ab, IgG and IgM antibody was calculated via S/CO multiplied by the maximum dilution factors.

##### Recombinant expression of SARS-CoV S protein, SARS-CoV-2 S protein and RBD

A gene encoding the ectodomain of a prefusion conformation-stabilized SARS-CoV-2 (GenBank: MN908947) S protein was synthesized, composed of SARS-COV-2 gene sequence (residue1-1208), a C-terminal T4 fibritin trimerization motif, an HRV3C protease and 8xHisTag. In addition, gene of SARS-CoV (GenBank: ABF65836) S protein comprised the same elements. To determine the blocking capacity of mAbs, we also constructed a gene comprising SARS-

COV-2 gene sequence (residue1-1208), a C-terminal T4 fibritin trimerization motif, an HRV3C protease, 8xHisTag and a -terminal Green fluorescent protein (mGamillus). Moreover, to express SARS-CoV-2 mutate and wildtype RBD, residues 319-541 fused to mouse IgG1 Fc domain. Recombinant expressions of these proteins were performed by the ExpiCHO™ expression system (Thermo Scientific, A29133). Briefly, plasmids encoding targeted proteins were transiently transfected into ExpiCHO cells by using ExpiFectamine™ CHO transfection kit (Thermo Scientific, A29129). The cell-free supernatants were obtained 5-7 days after transfection by centrifugation and filtration with a 0.22 μm filter. Subsequently, the proteins were purified by Ni Sepharose Excel resin, and stored in the PBS buffer.

**Specific memory B cell response and single B cell sorting**

RBD specific B cells were obtained in the same way as previously reported. PBMCs collected from 10 individuals were incubated with a cocktail containing live/dead-Aqua, CD3-PE-Cy7, CD19-BV786, CD27-BV650, anti-human IgM-PerCP-Cy5.5, anti-human IgG-BV421, RBD-FITC and biotinylated RBD, followed with Streptavidin-APC binding to biotinylated RBD. IgG+ memory B cells (CD19+CD3-CD27+ IgG+) that bind to RBD were single cell sorted from PBMC samples from each donor. Single cells were sorted by fluorescence activated cell sorting on an Aria III sorter (BD Biosciences) into 96-well PCR plates containing 20 μL per well of lysis buffer [5 μL of 5×first strand buffer (Invitrogen), 1.25 μL dithiothreitol (Invitrogen), 0.5 μL RNase Out (Invitrogen), 0.0625 μL Igepal (Sigma)]. Plates were stored at -80 °C prior to reverse transcription reaction.

**Single B cell PCR, cloning and expression of antibody**

Antibody variable genes (IgH, Igλ and Igκ) were amplified by RT-PCR and nested PCR reactions as previously described (*26*). The paired heavy and light chains were then cloned into expression vectors containing the constant regions of human IgG1 and light chain. The paired heavy and light chain expression cassettes were then transiently co-transfected into ExpiCHO cells with equal amounts of plasmids according to the manufacturer's instructions (Life Technologies), and antibodies were purified from culture supernatant 5-7 days after transfection, using a recombinant protein-A column (GE Healthcare).

**Antibody germline usage and phylogenetic analysis**

Antibody gene repertoire was analyzed for the variable region of IgG heavy and light chains using the IMGT V-quest webserver (http://www.imgt.org/IMGT/vquest). Phylogenetic analysis of antibody gene was performed by ggtree R package (*27*).

**Binding activity assay**

The binding activity of the mAbs against SARS-CoV-2 S protein were determined using an indirect ELISA. The mAbs were added to antigen-coated microwell plates, and incubated at 37°C for 30 min. Then, incubation of HRP-conjugated anti-human antibody at 37°C for 30 min to detect the bound mAbs, followed by washing five times. Finally, substrate solution was added to the wells for 15 min at 37 °C, and reaction was stopped by adding 50 µL of 2 M H2SO4. The optical density (OD) was measured at 450 nm with a reference wavelength of 630 nm. In addition, binding activity of the mAbs against SARS-CoV-2 RBD and SARS-CoV-1 S protein were determined by same method.

To determine the critical residues, mAbs were conjugated with horse radish peroxidase (HRP). Microwell plates were pre-coated with mutate RBD and wildtype RBD at 100 ng per well. mAbs-HRP were added at selected dilutions, at which OD readings was ~1.5 for wildtype RBD, and incubated at 37°C for 30 min followed by washing five times. Substrate solution was incubated for 15 min at 37 °C, and stopped by 50 µL of 2 M H2SO4. OD was determined at 450 nm with a reference wavelength of 630 nm. The reduction of binding activity of mAbs against mutated RBD was showed by comparation with OD against wildtype RBD.

**Blocking capacity of mAbs against SARS-CoV-2 S protein**

For SARS-CoV-2 S protein-blocking assay, mAbs were pre-made as 2-fold serial dilutions using DMEM containing 10% FBS. Aliquots (44 μL per well) of diluted samples and S protein probes (11 μL per well) were mixed in a 96-well plate with U shaped bottom. Half of the culture medium (50 μL) of 293T-ACE2iRb3 cell plate were gently removed, and 50 μL of sample/probe mixtures were added to each well. Cell image acquisitions performed with Opera Phenix (green, red and near-infrared channels in confocal mode) using a 20x water immersion objective at 1-hour after probe incubation in wash-free and live-cell conditions.

All quantitative image analyses were based on images that acquired by Opera Phenix. All image data were transfer to Columbus system (version 2.5.0, PerkinElmer Inc) for analysis. Multiparametric image analysis was performed as described in the following. The signals of blue channel or near-infrared channel were used to detect the nucleus. As the ACE2 is a membrane protein, the signals of ACE2-mRuby3 (red channel) were used to determine the cell boundary. Then, the cells were further segment into the regions of membrane (outer border: 0%, inner border: 15%), cytoplasm (outer border: 20%, inner border: 45%), and nucleus (outer border: 55%, inner border: 100%). The MFI of probe channel (Ex488/Em525) in the cytoplasmic region (cMFI). The MFI of ACE2-mRuby3 (Ex561/Em590) on the membrane were also calculated for inter-well normalization. The cMFI inhibition ratio (%) of the test sample was calculated using the following equation: [(cMFIpc-cMFItst)/(cMFIpc-cMFIblk)]×100%. In this formula, the cMFIpc is the cMFI value of probe-only well (as positive control), the cMFItst is the cMFI value of test well and the cMFIblk is the cMFI value of cell-only well. For each plate, 5 replicates of probe-only well and 1 cell-only well were included. The blocking capacity of mAbs were expressed as IC50.

**Neutralization capacity of mAbs determined by SARS-CoV and SARS-CoV-2 pseudovirus**

In order to construct the SARS-CoV-2 pseudovirus using VSV carrying the SARS-CoV-2 spike protein, the spike gene was codon optimized for expression in human cells, and the spike gene of SARS-CoV-2 with 18 amino acids truncated at the C-terminal was cloned into the eukaryotic expression vector pCAG to obtain pcag-ncovsde18. The plasmid pCAG-nCoVSde18 was transfected into Vero-E6. VSVdG-EGFP-G（Addgene，31842） virus was inoculated into cells expressing SARS-CoV-2 Sde18 truncated protein and incubated for 1 hour. Then the VSVdG-EGFP-G virus was removed from the supernatant and anti-VSV-G rat serum was added to block the remaining VSVdG-EGFP-G infection. The progeny virus will carry SARS-CoV-2 Sde18 truncated protein. After VSVdG-EGFP-G infection, supernatant was collected, centrifuged and filtered (Millipore, SLHP033RB) to obtain the SARS-CoV-2 pseudovirus without debris. SARS-CoV pseudovirus was constructed by the same method. Finally, pseudovirus was stored for use at -80°C.

To determine the neutralizing capacity, mAbs with 2-fold serial dilutions with 10% FBS-DMEM from 2 μg/mL were mixed with diluted SARS-CoV and SARS-CoV-2 pseudovirus (MOI = 0.05), incubated at 37°C for 1hour. A mixture of 80 μL was added to the precoated BHK21-hACE2 cells. After incubation for 12 hours, post-infection cells were fluorescently imaged using Opera phenix or Operetta CLS (Perkinelmer), and quantitatively analyzed by Columbus image management analysis software to detect the number of green fluorescent positive cells. The inhibition rate was calculated by reduction of GFP positive cells with presence of mAbs compared with the untreated control wells.

**Competition binding assay for neutralizing antibodies by ELISA and cluster analysis**

Briefly, the unlabeled mAbs (50 μg per well) or PBS were added to RBD-coated 96-well microplates and then incubated for 30 min at 37 °C. Next, HRP-conjugated mAbs were added at selected dilutions, at which OD readings was ~1.5 with PBS present. After incubation for 30 min at 37 °C, the microplates were rinsed and the color was developed. The competitive ability was measured quantitatively by comparing OD in the presence or absence of competitor mAbs, and transformed using the formula log2 ($OD_{inhibited}$/ $OD_{original}$). For mAbs to be clustered by competitive ability, clustering distance was calculated by Euclidean, and cluster by ward.D2 method, using pheatmap R package (version: 1.0.12).

**Statistical analysis**

To compare continuous variables, the Mann-Whitney U test and non-paired t test were performed. Linear regression model and Spearman test were used for correlation analyses. For difference analysis, p values less than 0.05 are considered statistically significant. GraphPad Prism version 8.0.1 was used for all statistical calculations.

**Fig. S1. Characteristics analysis of humoral immune response by COVID-19 convalescent plasma. (A)** Correlation between days after symptom onset and plasma antibody titer including anti-RBD antibody, anti-RBD IgG and anti-RBD IgM, using Spearman correlation test. **(B)** Correlation test between anti-RBD titers and PSV SARS-CoV-2 neutralizing capacity are determined by Spearman correlation test. r and P values of the correlation are indicated.

**Fig. S2. Identification of SARS-CoV-2 RBD-specific memory B cells and isolation of SARS-CoV-2 RBD-specific antibodies. (A)** SARS-CoV-2 RBD-specific memory B cells are identified as CD3-/CD19+/CD27+/SARS-CoV-2 RBD+, and the percentage of RBD-specific B cells is indicated. **(B and C)** The BCR (B cell receptor) subtypes of RBD-specific memory B cells are analyzed by goat anti-human IgG and goat anti-human IgM, then are statistically analyzed. **(D)** Recombinant monoclonal antibodies with SARS-CoV-2 RBD specificity are identified by ELISA. Gray line indicates limitation of anti-RBD antibodies detection.

**Fig. S3. Phylogenetic analysis of heavy chain gene of SARS-CoV-2 RBD-specific mAbs.** Maximum-likelihood phylogenetic tree of fully heavy chain of RBD-specific antibodies (N=77). Each color represents heavy chain sequence of SARS-CoV-2 RBD-specific mAbs from different convalescent individuals.

**Fig. S4. Gene repertoire analysis of SARS-CoV-2 RBD-specific mAbs.** V gene frequencies for heavy chain **(A)** and light chain **(B)** of SARS-CoV-2 RBD-specific antibodies. Colors indicate different convalescent individuals. Germline of VH are determined using the Immunogenetics (IMGT).

**Fig. S5. Binding activity to S protein and neutralizing capacity against SARS-CoV-2 pseudovirus of SARS-CoV-2 RBD-specific mAbs. (A and B)** Binding activity of mAbs to SARS-CoV S protein are compared among individual in B, and to SARS-CoV-2 S protein in C. Black line indicates mean value of $EC_{50}$. **(C and D)** Neutralizing capacity of mAbs against SARS-CoV-2 are compared among individual in **C**, and blocking capacity of mAbs in **D**.

Neutralizing capacity are tested by SARS-CoV-2 pseudovirus. Blocking assay is performed by incubating mixture of antibodies and SARS-CoV-2 S protein with ACE2-expressing cells. Black line indicates mean $IC_{50}$.

**Fig. S6. Neutralization of SARS-CoV-2 RBD-specific mAbs against SARS-CoV-2 pseudovirus.** Red indicates mAbs obtained from P03 convalescent individual.

**Fig. S7. The correlation between neutralization potency and blocking capability of SARS-CoV-2 RBD-specific mAbs.** The scatter plot depicting neutralizing capacity and blocking capacity of specific mAbs from different individuals annotated by colors.

**Fig. S8. Analysis of CDRH3 length of antibodies derived from VH 3-53/66.** **(A)** Repertoire information of RBD-specific antibodies composed of VH 3-53/66. **(B)** Length distribution of CDRH3 for RBD-specific antibodies derived from VH 3-53/66 by comparison with the remaining VH germline encoding antibodies. **(C)** Correlation of CDRH3 length and binding activity to SARS-CoV-2 S protein is performed for specific antibodies derived from of VH 3-53/66.

**Fig. S9.** Correlation analysis for days after symptom onset and mean binding activity of SARS-CoV-2 RBD-specific mAbs from corresponding convalescent individuals using Spearman correlation test. r and P values of the correlation are indicated.

**Fig. S10. Competition ELISA for neutralizing mAbs.** Competition ELISA is performed by using naked mAbs to block HRP-coupled mAbs, and ELISA signal for each HRP-coupled mAb is normalized to the signal in the absence of naked mAbs. The heat map of competition ELISA data is shown, with parameters colored continuously from white (0, corresponding to 0% inhibition) to red (4, corresponding to 93.7% inhibition) in the scale bar.

**Fig. S11. Epitope mapping of mAbs by clustering analysis and functional characterization.** By competition ELISA data, neutralizing mAbs are clustered into 6 group, Cluster1-6, and corresponding epitopes to each mAb cluster are defined as Site1-6. The color ranging from red to blue represented blocking potency against other antibodies (4.321 corresponding to 95% blocking rate and 0.074 corresponding to 5% blocking rate). The source of information and neutralization potency of each mAb are also indicated by different colors.

**Fig. S12. Analysis of blocking capability against SARS-CoV-2 S protein binding to ACE2. (A)** Blocking capacity of NAbs targeting sites S2-6 are compared with that of NAbs recognizing S1. **(B)** The neutralizing capacity and blocking capacity of NAbs recognizing site S4 are analyzed, and NAbs ID are indicated in figure.

**Fig.S13. Identification of SARS-CoV-2 RBD critical residues recognized by NAbs using selected amino acid substitution. (A)** Mutate residues of SARS-CoV-2 RBD shown in pink. **(B)** Mutation of residues leading to damaging effect on SARS-CoV-2 RBD activity. Dash line indicates 25% binding activity of mutant RBD relative to wild type RBD. **(C)** The selected amino acid of RBD are mutated to alanine or arginine on purpose. Binding activity of sites S1-6 representative mAbs to wild-type (WT) and mutant RBD was measured by ELISA. The binding capacity to mutate RBD is normalized by binding to wild type RBD. Lines denote 10% binding activity relative to wild type RBD and 25% binding activity relative to wild type RBD. Residues reducing binding activity by more than 75% are identified critical residues for representative NAbs.

**Fig. S14. Identification of sites S1-6 spatial position. (A)** Structure of the RBD highlighting the critical residues interfering binding activity of representative NAbs, red denotes residues reducing binding activity by more than 75%. **(B)** Conservative analysis of sites S1-6, carmine denotes different residues between SARS-CoV-2 RBD and SARS-CoV RBD.

**Fig. S15.** CDRH1-3 sequence analysis of mAbs derived from IGHV3-53/66, including P05-5C4 derived from IGHV3-53 and P02-3C11 derived from IGHV3-66 targeting site S1.

#### Fig. S1

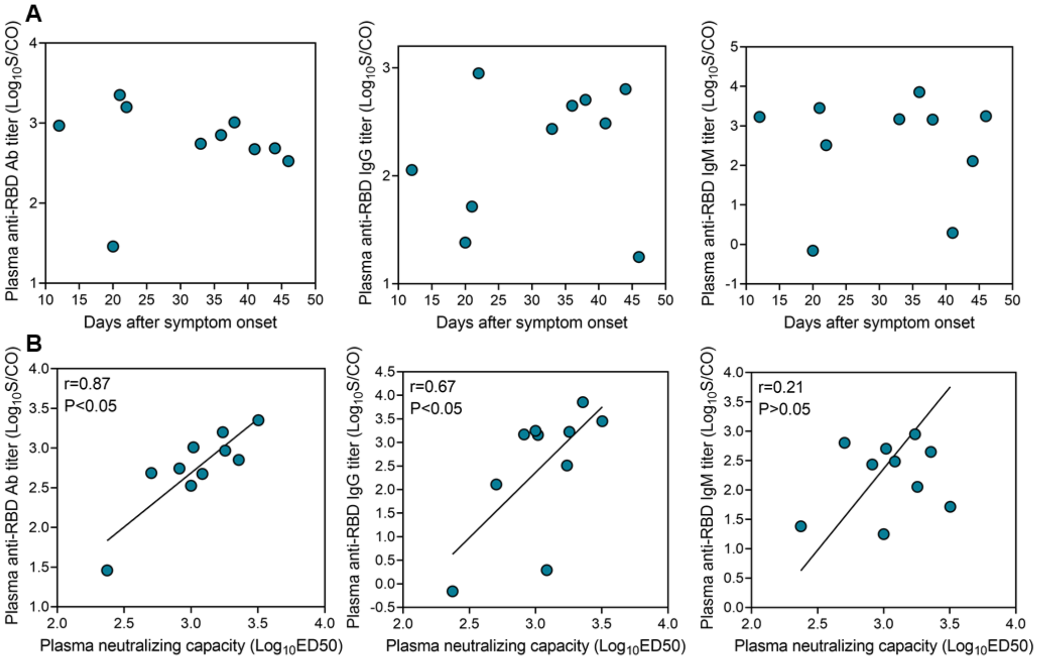
A
Plasma anti-RBD Ab titer (Log10S/CO)
Days after symptom onset
Plasma anti-RBD IgG titer (Log10S/CO)
Days after symptom onset
Plasma anti-RBD IgM titer (Log10S/CO)
Days after symptom onset
B
r=0.87
P<0.05
Plasma anti-RBD Ab titer (Log10S/CO)
Plasma neutralizing capacity (Log10ED50)
r=0.67
P<0.05
Plasma anti-RBD IgG titer (Log10S/CO)
Plasma neutralizing capacity (Log10ED50)
r=0.21
P>0.05
Plasma anti-RBD IgM titer (Log10S/CO)
Plasma neutralizing capacity (Log10ED50)

**Fig. S2**

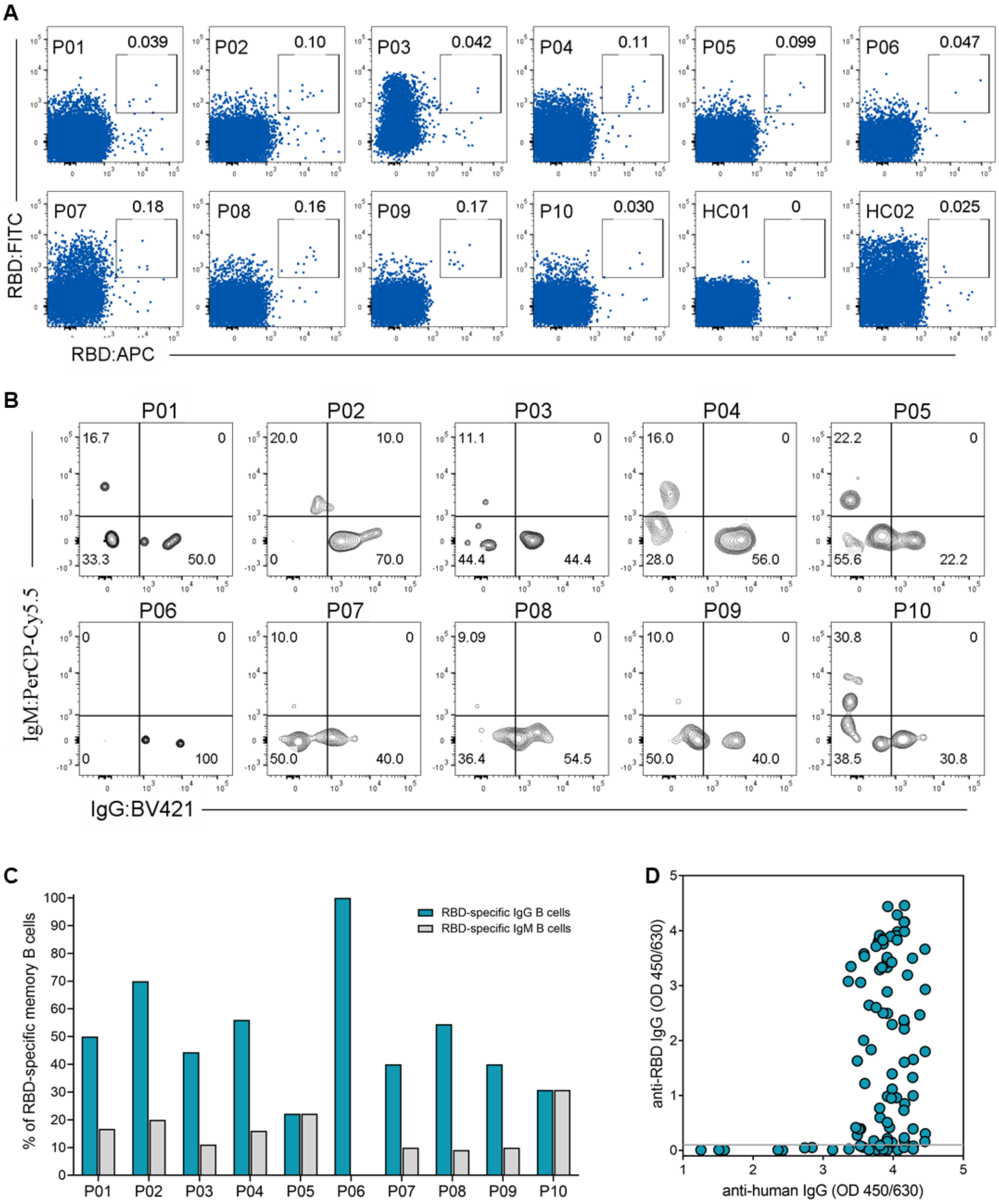

A
RBD:FITC
RBD:APC
P01 0.039
P02 0.10
P03 0.042
P04 0.11
P05 0.099
P06 0.047
P07 0.18
P08 0.16
P09 0.17
P10 0.030
HC01 0
HC02 0.025
B
IgM:PerCP-Cy5.5
IgG:BV421
P01 16.7 0 33.3 50.0
P02 20.0 10.0 0 70.0
P03 11.1 0 44.4 44.4
P04 16.0 0 28.0 56.0
P05 22.2 0 55.6 22.2
P06 0 0 0 100
P07 10.0 0 50.0 40.0
P08 9.09 0 36.4 54.5
P09 10.0 0 50.0 40.0
P10 30.8 0 38.5 30.8
C
% of RBD-specific memory B cells
RBD-specific IgG B cells
RBD-specific IgM B cells
P01 P02 P03 P04 P05 P06 P07 P08 P09 P10
D
anti-RBD IgG (OD 450/630)
anti-human IgG (OD 450/630)

#### Fig. S3

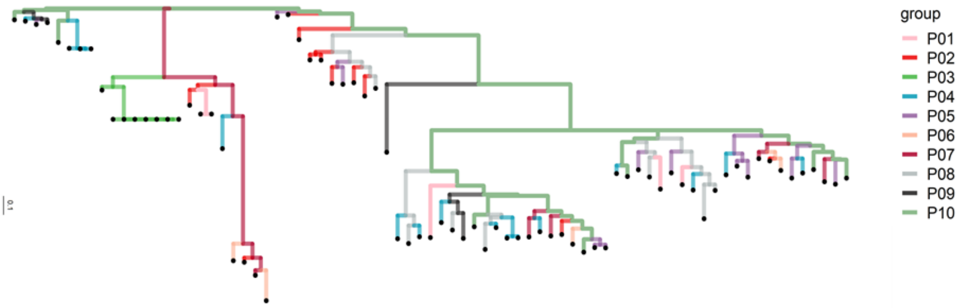


#### Fig. S4

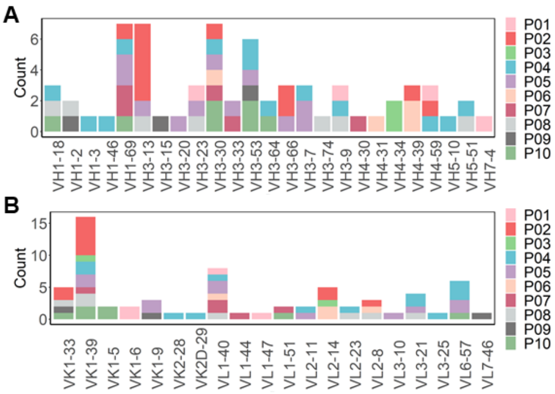


#### Fig. S5

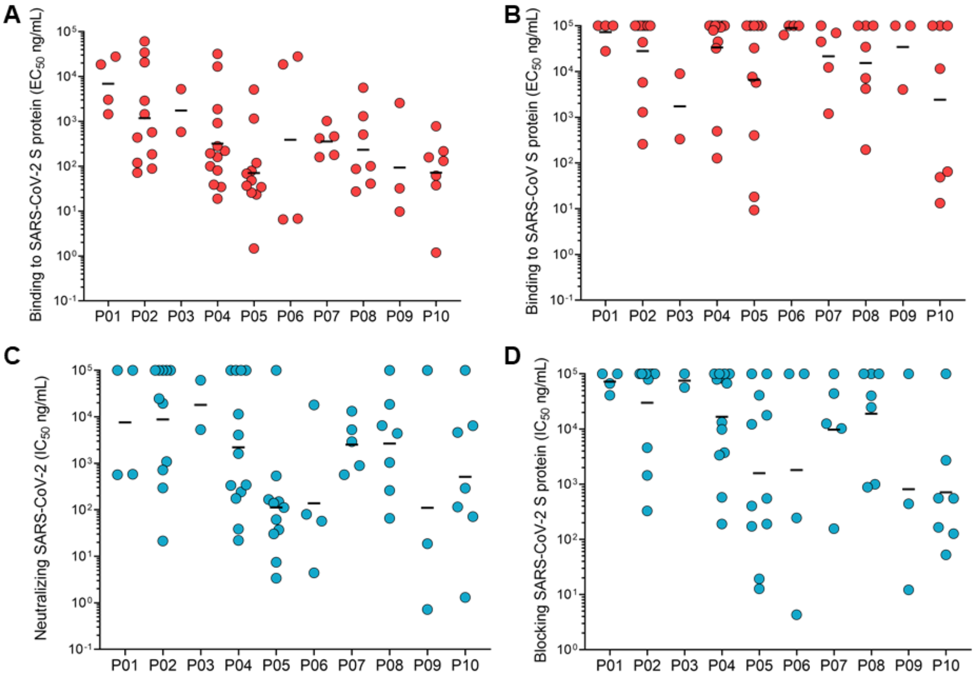

**Fig. S6**

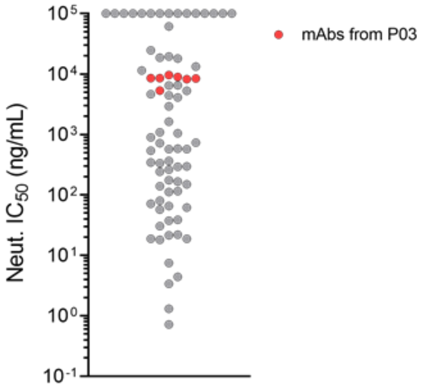


**Fig. S7**

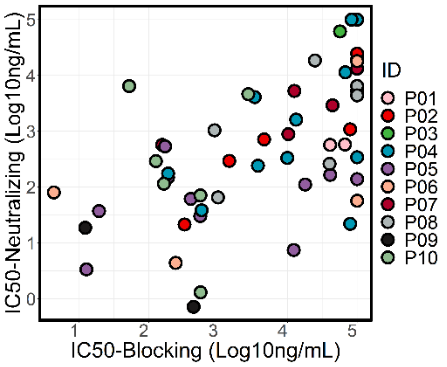


**Fig. S8**

**A**

| Donor | ID | V-GENE | J-GENE | D-GENE | CDRH3 | Length |
|---|---|---|---|---|---|---|
| P02 | 3C11 | IGHV3-66*01 | IGHJ6*02 | IGHD3-10*01 | ARGLVASYRYGMDV | 14 |
| P02 | 3D4 | IGHV3-66*01 | IGHJ4*02 | IGHD2-21*01 | ARDWGGRRFDY | 11 |
| P05 | 5A6 | IGHV3-66*01 | IGHJ4*02 | IGHD5-24*01 | ARDRGGDTFDY | 11 |
| P05 | 5C4 | IGHV3-53*04 | IGHJ6*02 | IGHD2-21*01 | ARDLVVYGMDV | 11 |
| P04 | 6B2 | IGHV3-53*01 | IGHJ6*02 | IGHD2-8*02 | ARSLEVYGLDV | 11 |
| P04 | 6D4 | IGHV3-53*01 | IGHJ4*02 | IGHD5-12*01 | ARDRAWGGYDWTFDY | 15 |
| P10 | 6F11 | IGHV3-53*01 | IGHJ4*02 | IGHD6-13*01 | ARLPYSSNWES | 11 |
| P10 | 6G5 | IGHV3-53*04 | IGHJ3*02 | IGHD3-10*01 | ARVVSDAFDI | 10 |
| P09 | 6G8 | IGHV3-53*01 | IGHJ6*02 | IGHD1-1*01 | ARDLDYYGMDV | 11 |

**B**

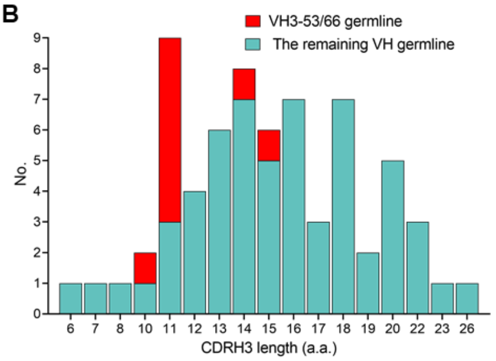


**C**

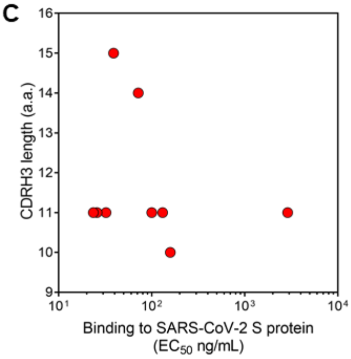

**Fig. S9**

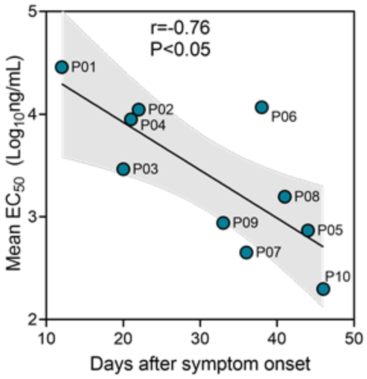


**Fig. S10**

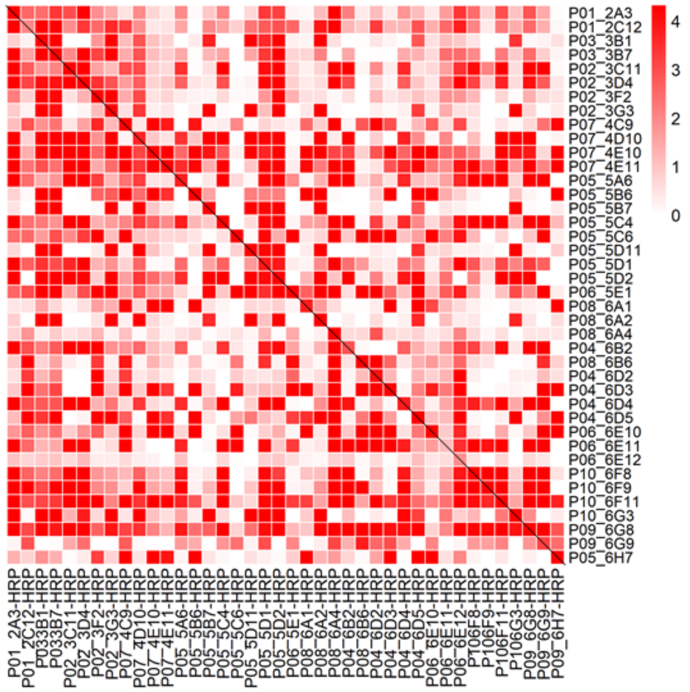

#### Fig. S11

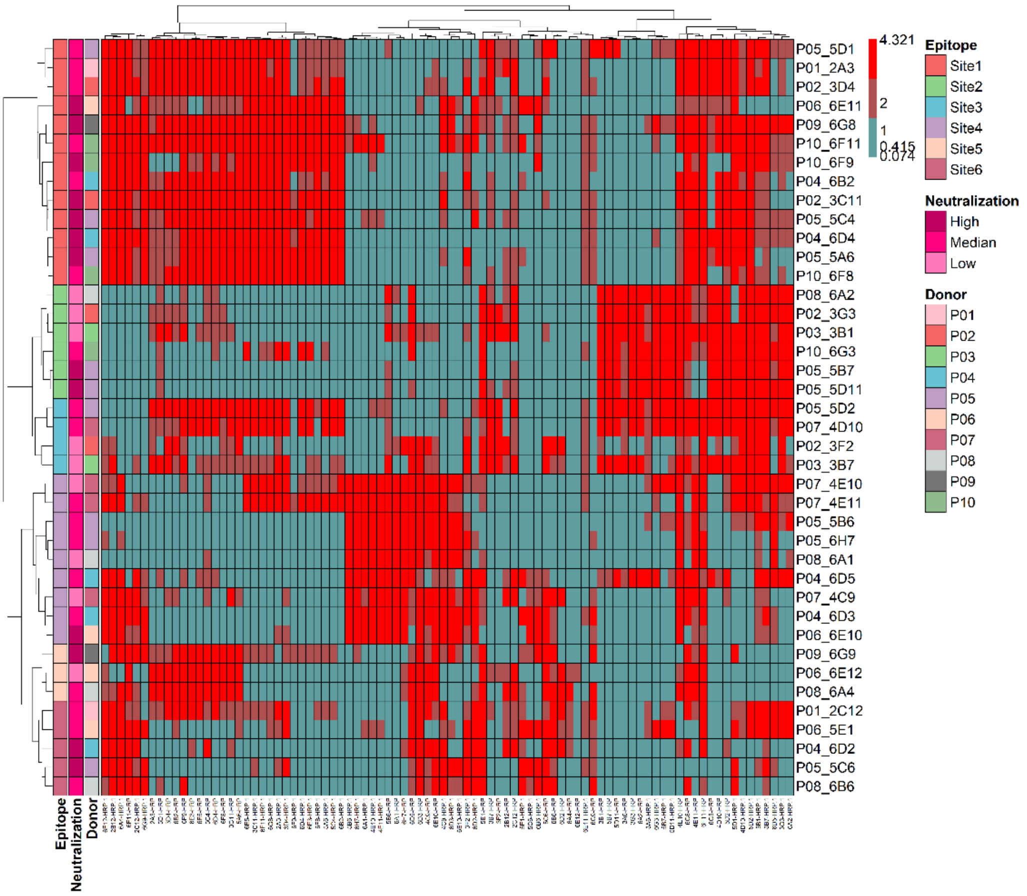


#### Fig. S12

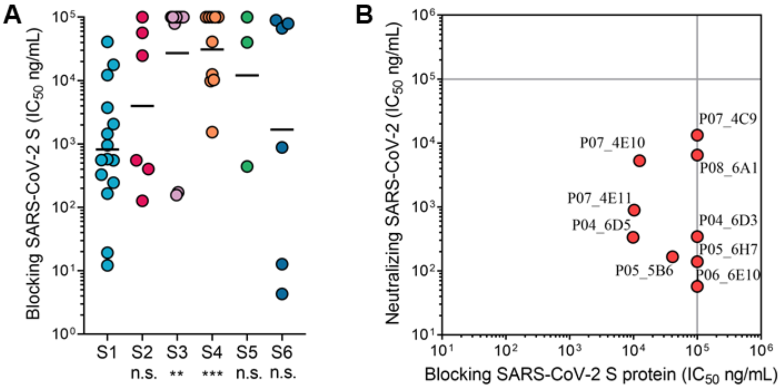

#### Fig. S13

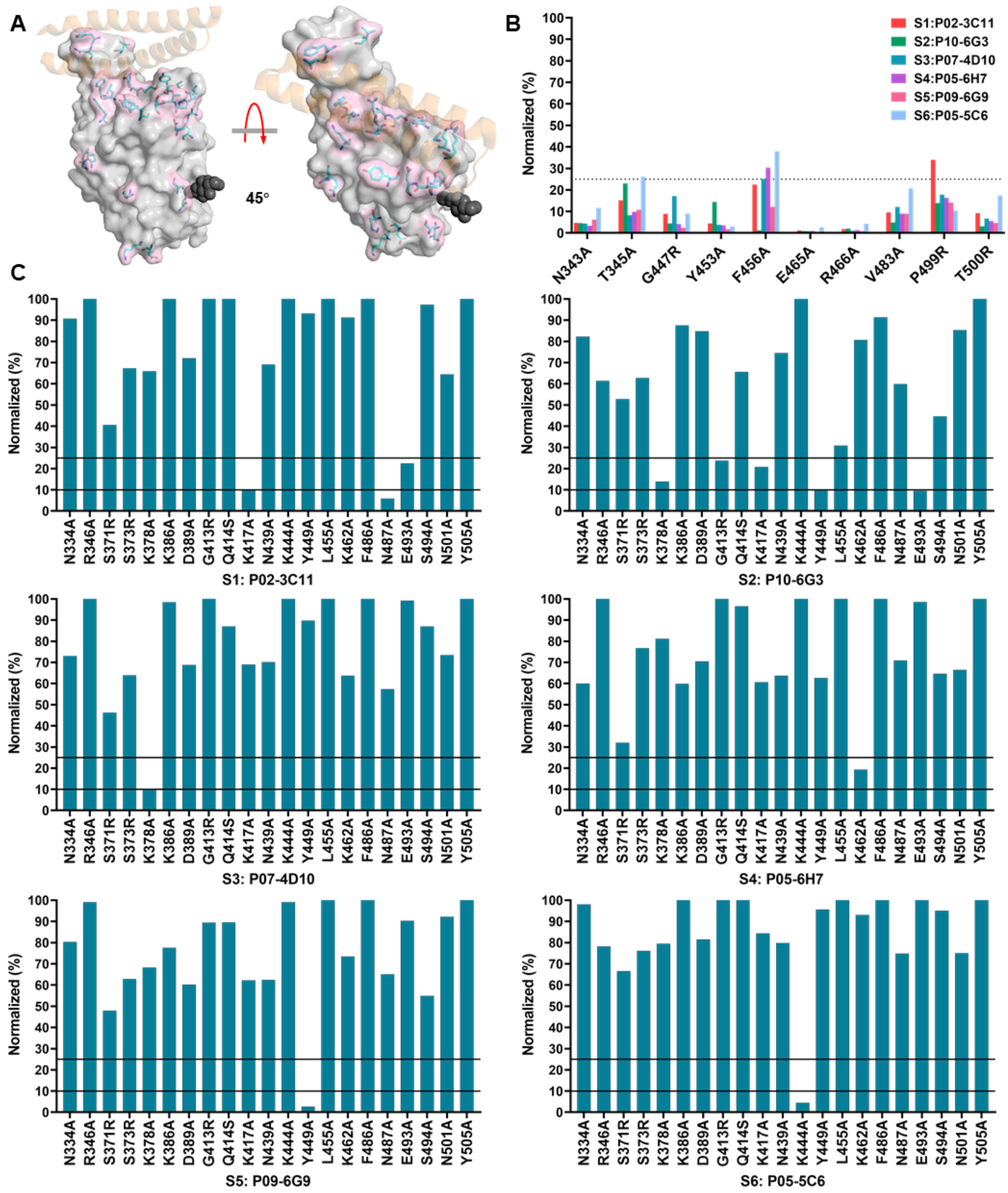
A
45°
B
S1:P02-3C11
S2:P10-6G3
S3:P07-4D10
S4:P05-6H7
S5:P09-6G9
S6:P05-5C6
Normalized (%)
N343A
T345A
G447R
Y453A
F456A
E465A
R466A
V483A
P499R
T500R
C
S1: P02-3C11
S2: P10-6G3
S3: P07-4D10
S4: P05-6H7
S5: P09-6G9
S6: P05-5C6
N334A
R346A
S371R
S373R
K378A
K386A
D389A
G413R
Q414S
K417A
N439A
K444A
Y449A
L455A
K462A
F486A
N487A
E493A
S494A
N501A
Y505A

#### Fig. S14

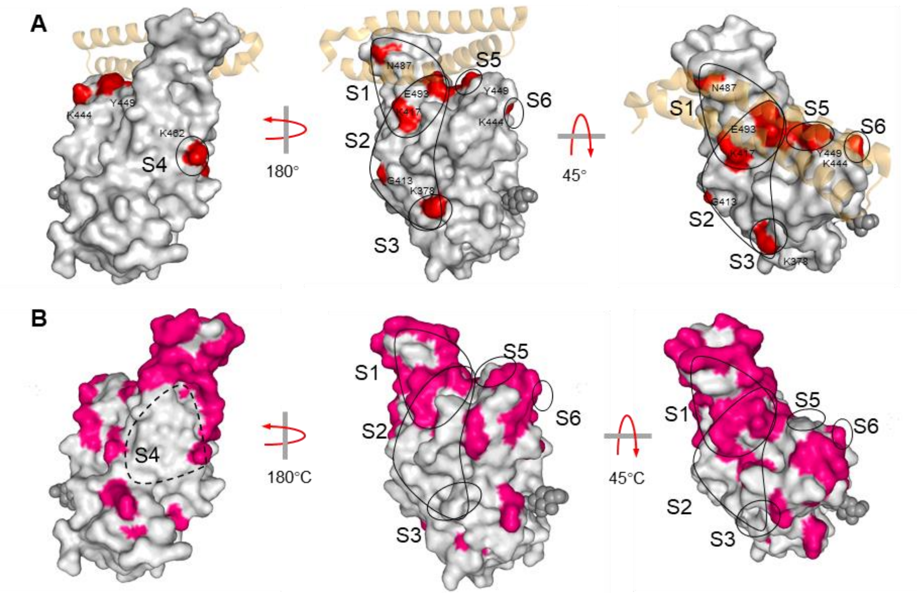


#### Fig. S15

| CDRH1 | $G_{26}$ | $F_{27}$ | $I_{28}$ | $V_{29}$ | $S_{30}$ | $S_{31}$ | $N_{32}$ | $Y_{33}$ |
|---|---|---|---|---|---|---|---|---|
| C102 | - | - | - | - | - | - | - | - |
| CC12.1 | - | L | T | - | - | - | - | - |
| CC12.3 | - | - | T | - | - | - | - | - |
| B38 | - | - | - | - | - | - | - | - |
| C105 | - | - | T | - | - | - | - | - |
| P05-5C4 | - | - | T | - | - | - | - | - |
| P02-3C11 | - | - | T | - | - | R | - | - |

| CDRH2 | $I_{51}$ | $Y_{52}$ | $S_{53}$ | $G_{54}$ | $G_{55}$ | $S_{56}$ | $T_{57}$ |
|---|---|---|---|---|---|---|---|
| C102 | - | - | - | - | - | - | - |
| CC12.1 | - | - | - | - | - | - | - |
| CC12.3 | - | - | - | - | - | - | - |
| B38 | - | - | - | - | - | - | - |
| C105 | - | - | - | - | - | - | - |
| P05-5C4 | - | - | - | - | - | - | - |
| P02-3C11 | - | - | - | - | - | - | - |

| CDRH3 | a.a. | Length |
|---|---|---|
| C102 | ARDYGDYYFDY | 11 |
| CC12.1 | ARDLDVYGLDV | 11 |
| CC12.3 | ARDFGDFYFDY | 11 |
| B38 | AREAYGMDV | 9 |
| C105 | ARGEGWELPYDY | 12 |
| P05-5C4 | ARDLVVYGMDV | 11 |
| P02-3C11 | ARGLVASYRYGMDV | 14 |

**Table. S1. Information of COVID-19 convalescent individuals**. F: female M: male.

| ID | Age | Gender | Severity | Symptom onset date | Hospitalization date | PBMC collection date | First Symptoms | Chronic basic disease | Infection of other virus | CT | SARS-CoV-2 RNA |
|---|---|---|---|---|---|---|---|---|---|---|---|
| P01 | 54 | F | mild | 1.25 | 1.26-2.11 | 2.6 | Cough | hypertension | none | Bilateral | + |
| P02 | 64 | F | mild | 2.7 | 2.8-2.24 | 2.29 | Fever | none | none | Bilateral | + |
| P03 | 69 | M | mild | 2.9 | 2.9-2.25 | 2.29 | Fever | hypertension, hematencephalon | none | Bilateral | + |
| P04 | 33 | F | mild | 2.8 | 2.9-2.26 | 2.29 | Fever | none | HBV | unilateral | + |
| P05 | 40 | M | mild | 1.21 | 1.26-2.20 | 3.5 | Fever | none | none | Bilateral | + |
| P06 | 68 | M | mild | 1.28 | 1.30-2.20 | 3.6 | Fever | hypertension | none | Bilateral | + |
| P07 | 50 | F | mild | 1.3 | 2.4-2.20 | 3.7 | Fever, Cough | none | none | Bilateral | + |
| P08 | 47 | M | mild | 1.26 | 1.28-2.20 | 3.7 | Fever | none | HBV | Bilateral | + |
| P09 | 71 | M | mild | 2.5 | 2.7-2.23 | 3.9 | Fever, Cough | hypertension, diabetes | none | Bilateral | + |
| P10 | 44 | M | mild | 1.28 | 1.30-2.27 | 3.14 | Fever, Cough | diabetes | none | Bilateral | + |

**Table. S2. Plasma anti-RBD antibody titers and neutralization capacity for COVID-19 convalescent individuals.**

| ID | Days after symptom onset | Anti-RBD antibodies titer (S/CO) | Anti-RBD IgG titer (S/CO) | Anti-RBD IgM titer (S/CO) | Neutralization capacity (ED50) |
|---|---|---|---|---|---|
| P01 | 12 | 931.58 | 113.16 | 1685.71 | 1798.00 |
| P02 | 22 | 1584.21 | 889.47 | 324.76 | 1728.00 |
| P03 | 20 | 28.74 | 24.05 | 0.69 | 235.90 |
| P04 | 21 | 2247.37 | 51.89 | 2819.05 | 3185.00 |
| P05 | 44 | 484.21 | 634.21 | 128.57 | 505.50 |
| P06 | 38 | 1021.05 | 505.26 | 1447.62 | 1041.00 |
| P07 | 36 | 710.53 | 445.26 | 7152.38 | 2276.00 |
| P08 | 41 | 473.68 | 305.79 | 1.95 | 1215.00 |
| P09 | 33 | 552.60 | 272.10 | 1476.20 | 820.00 |
| P10 | 46 | 334.70 | 17.70 | 1761.90 | 1001.00 |
